# Genetic diversity in horseshoe bat ACE2 and sarbecovirus spike proteins mutually shape one another

**DOI:** 10.1101/2025.10.26.684670

**Authors:** Wenye Li, Shoma Sato, Taichiro Takemura, Xiao Niu, Daniel Arnold, Maximillian Stanley Yo, Jarel Elgin Tolentino, Shigeru Fujita, Yusuke Kosugi, Nguyen Thi Thu Thuy, Le Thi Quynh Mai, Nguyen Thanh Luong, Hidetaka S. Oshima, Kanata Matsumoto, Kazuhiro Sawada, Hiroaki Akasaka, Wataru Shihoya, Jumpei Ito, Yunlong Cao, Futoshi Hasebe, Vu Dinh Thong, Osamu Nureki, Spyros Lytras, Kei Sato

## Abstract

Angiotensin-converting enzyme 2 (ACE2) serves as the entry receptor for a wide diversity of sarbecoviruses naturally harboured by horseshoe bats (genus *Rhinolophus*). Despite the extensive circulation of these viruses in many horseshoe bat species, the potential interactions between virus and receptor evolution remain poorly understood. We sampled individuals of the intermediate horseshoe bat (*Rhinolophus affinis*) across Vietnam and identified 15 genotypes of ACE2 proteins, 10 of which are previously unreported. Phylogenetic analysis and infectivity assays with a panel of 36 sarbecovirus spike proteins indicated that the *R. affinis* ACE2 phylogeny has geographic structuring and genotypes originating from different geographic regions exhibit distinct infectivity phenotypes. We detected site-specific positive selection on ACE2 site 24 with the associated substitutions largely affecting the receptor’s sarbecovirus infectivity profile. Together, our findings suggest that the *R. affinis* within-species ACE2 diversity has likely been shaped through selection by past sarbecovirus infection. Similarly on the virus end, we use mutagenesis assays and structural analysis through cryo-EM, to delineate the proximal evolution of the Ra22QT77 spike defined by specialization to the ACE2 genotypes of horseshoe bats found in and near southern Vietnam, where the virus was sampled. Our findings contribute to a better understanding of host-sarbecovirus co-evolution dynamics and provide valuable insights into the receptor usage determinants of these viruses.

**Highlights:**

- We identify a total of 15 ACE2 genotypes in *R. affinis* bats from Vietnam.
- ACE2 intraspecific polymorphism is geographically separated and associated with distinct sarbecovirus infectivity.
- Site 24 of *R. affinis* ACE2 experiences positive selection and controls susceptibility to sarbecoviruses.
- Bat ACE2s and sarbecovirus spikes are bidirectionally shaped by each other’s evolution.

## Introduction

Coronaviruses (CoV, family *Coronaviridae*) are enveloped, positive-sense, single-stranded RNA viruses possessing some of the largest RNA genomes of ∼26-32 kb^1^. The subfamily *Orthocoronavirinae* is taxonomically divided into four genera: *Alphacoronavirus*, *Betacoronavirus*, *Gammacoronavirus*, and *Deltacoronavirus*^2^. Alphacoronaviruses and betacoronaviruses only infect mammals, whereas gammacoronaviruses and deltacoronaviruses circulate mainly in birds, with few known mammal-infecting members^3,4^. Over the past two decades, three highly pathogenic betacoronaviruses have caused major human outbreaks: the severe acute respiratory syndrome CoV (SARS-CoV; 2002-2004)^5,6^, the Middle East respiratory syndrome coronavirus (MERS-CoV, first detected in 2012 and still causing sporadic zoonotic infections)^7^, and the severe acute respiratory syndrome coronavirus 2 (SARS-CoV-2, responsible for the COVID-19 pandemic beginning at the end of 2019)^8,9^. Phylogenetic and ecological evidence indicates that SARS-CoV and SARS-CoV-2 originated in horseshoe bats and possibly reached humans via intermediate hosts, underscoring the zoonotic potential of betacoronaviruses and the need for sustained bat surveillance^10^.

Bats are widely recognized as natural reservoirs of diverse zoonotic pathogens, particularly betacoronaviruses. Horseshoe bats (family *Rhinolophidae*) are known to harbour a wide diversity of SARS-related betacoronaviruses, exhibiting high positivity in the wild^10,11^. The genus *Rhinolophus* comprises of more than 100 extant species and is split into two lineages: Afro-Palaearctic and Asian, which diverged roughly 17-27 million years ago, providing a long evolutionary window for reciprocal diversification of these hosts and the viruses by which they are infected^12^. The Chinese rufous horseshoe bat (*R. sinicus*) was the first species shown to harbour SARS-related coronaviruses (SARSr-CoVs) that are the closest known relatives to SARS-CoV, establishing this species as the likely wildlife source of the virus^13,14^. Since 2021 several additional sarbecoviruses sharing more than 95% genome identity with SARS-CoV-2 have been reported from other horseshoe bat species: BANAL-20-52, BANAL-20-103 and BANAL-20-236 from *R. malayanus*, *R. pusillus* and *R. marshalli* in Laos respectively, and RaTG13 from *R. affinis* in Yunnan, China^9,15,16^. All these sarbecoviruses use angiotensin-converting enzyme 2 (ACE2) as their entry receptor and are capable of utilising their natural bat hosts as well as the human orthologues of the receptor^15,17^.

ACE2 is a protein with multiple functions that counter-balances classical renin-angiotensin signalling and serves as the primary cellular receptor for most sarbecoviruses^18,19^. Functional studies have confirmed that orthologues from multiple bat genera, including *Rhinolophus*, *Hipposideros* and *Miniopterus,* can mediate entry of phylogenetically diverse sarbecoviruses^20–22^. One species example is the *Rhinolophus affinis* ACE2, which supports infection by SARS-CoV-2, its close bat relatives RaTG13 and BANAL-20-52/-236, as well as a pangolin CoV, PCoV-GD/1/2019, underscoring the remarkable plasticity of the spike-ACE2 interface and the capacity of sarbecoviruses to exploit receptors from taxonomically distant hosts^15,21,23^.

Despite being generally conserved across many species, ACE2 displays pronounced intraspecific polymorphism in bats. Allelic diversity has been previously recorded in *Hipposideros armiger*, *R. sinicus*, *R. ferrumequinum* and *R. affinis*^24–26^. In *R. affinis* alone, 23 previously described nucleotide haplotypes encode seven protein variants (RA-01 to RA-07)^24^. Studies on the function and structure of these ACE2s reveal that substitutions at key interface residues 34, 38, and 83 can modulate RaTG13 spike pseudovirus entry by nearly 30-fold, whereas the same changes have minimal impact on SARS-CoV-2 and BANAL-20-52/-236, emphasizing pronounced virus-specific differences in receptor usage^24^. Such polymorphism provides multiple targets that sarbecoviruses can act on, potentially shaping the ecological landscape of host susceptibility.

How ACE2 diversity functions to promote the molecular arms race between bats and their sarbecoviruses currently remains poorly understood. Selection analysis of *R. sinicus* ACE2s reveals recurrent signal of positive selection, with most sites under selection coinciding with residues that directly interact with the viral spike^27,28^. On the virus side, deep-mutational scanning experiments demonstrate that ACE2 binding is a highly evolvable trait, since single amino acid substitutions can largely enhance or lower affinity to different ACE2 orthologues, expanding or contracting host range in a single mutational step^25,29^. Additionally, recombination analysis on bat sarbecovirus genomes reveals recurrent exchange of receptor-binding domains (RBDs), suggesting that this process contributes to spike diversification and geographic spread^30^.

Together, the mutational plasticity of the RBD, frequent recombination, and host-driven selection acting on ACE2 create a dynamic co-evolutionary landscape that both constrains and enables cross-species transmission. In this study, we aim to investigate the ACE2 polymorphism within the intermediate horseshoe bat, *R. affinis,* and examine how receptor diversity shapes the evolution of sarbecoviruses and the plausible molecular arms race between virus and host. We have systematically characterised new *R. affinis* ACE2 genotypes and quantified their binding and entry efficiency with a broad panel of representative sarbecovirus spikes. Our data show that single-residue polymorphisms in these ACE2s can modulate susceptibility to sarbecoviruses, highlight genotypes that restrict or facilitate infection by high-risk viral lineages, and provide evidence for virus-driven selection on the receptor within a single horseshoe bat species.

## Results

### Sample collection and species identification

Between 2021 and 2022, we captured a total of 245 *Rhinolophus* bat individuals from four distinct locations across Vietnam and retrieved organ samples including kidney, liver and intestines. Three of the sampling sites are located in northern Vietnam: Tam Dao, Lang Son, and Cat Ba (northern bat cohort); while the last site, Bach Ma, is located in southern Vietnam (southern bat cohort) **(Figure 1A)**. Notably, Cat Ba is an island in northern Vietnam. Through morphological characterization^31^ and genetic barcoding based on the cytochrome B (CYTB) gene^32^, a total of 155 sampled individuals were conclusively identified as *R. affinis*. Specifically, 38 were captured at Bach Ma and 20 at Tam Dao in 2021, followed by an additional 74 from Bach Ma, 19 from Lang Son, and 4 from Cat Ba in 2022 **(Table S1)**.

**Figure 1.**
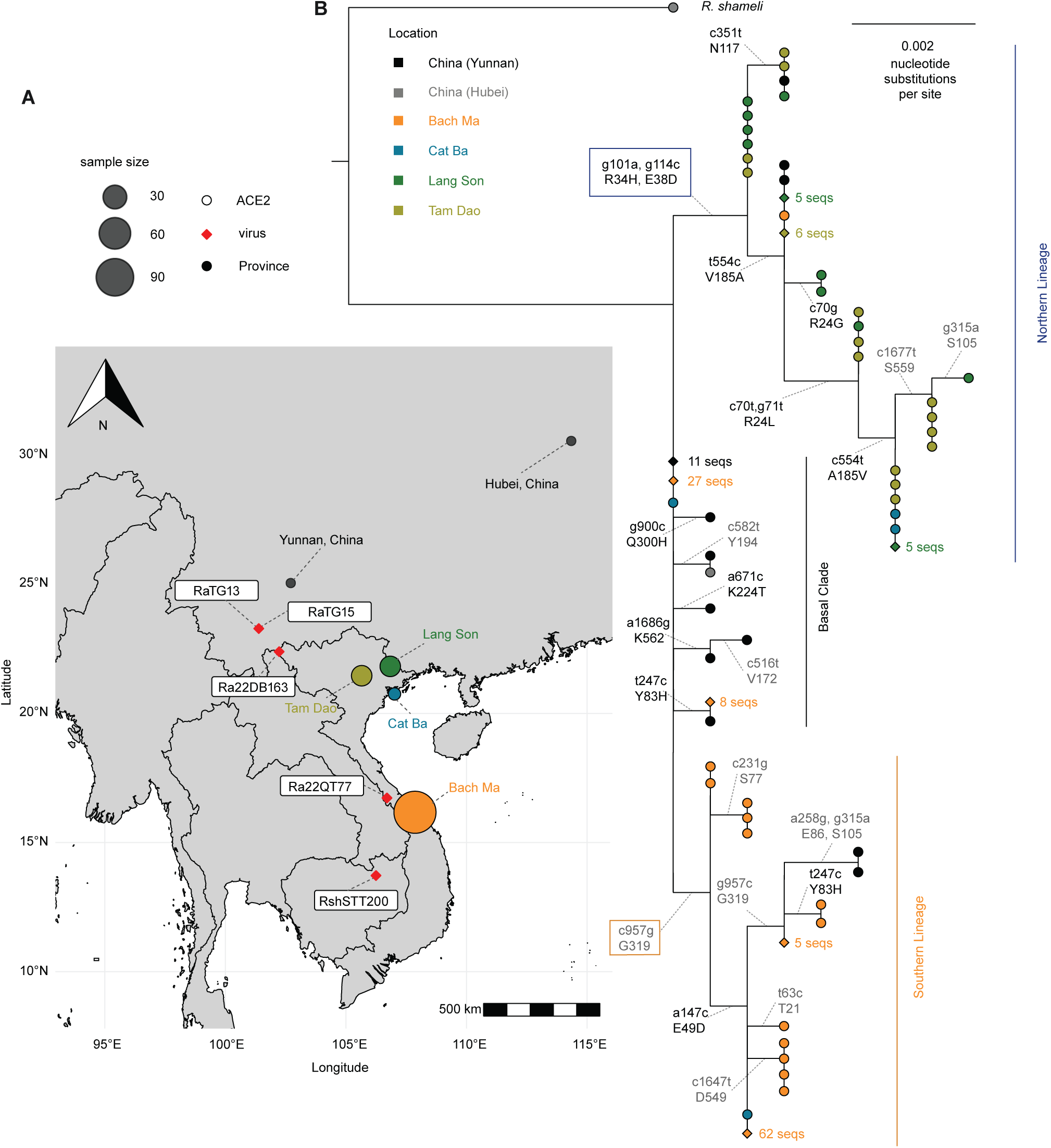
Geographic distribution and evolution of *R. affinis* ACE2. **(A)** Geographic distribution of *R. affinis* sampling in Vietnam with reference ACE2 genotypes and sarbecoviruses. A map of Vietnam within Southeast Asia is shown. Field-captured *R. affinis* individuals are plotted at their GPS coordinates **(Table S1)** as coloured dots; colors correspond to each sampling location and dot diameter to the number of samples at each location. Two provinces in China where ACE2 genotypes C-Ra-01 and C-Ra-02 were previously sampled in^24^ are marked as dark circles. The sampling locations of five sarbecoviruses (RaTG13, RaTG15, Ra22DB163, Ra22QT77, and RShSTT200, **Table S3**) are plotted as red squares at their reported sampling coordinates. **(B)** Maximum likelihood tree of the 182 *R. affinis* ACE2 coding sequences. A total of 159 identified sequences and 23 publicly available sequences are included. The phylogeny constructed based on only the left-hand partition of the alignment is shown. The K2P+R2 substitution model was used for the phylogeny. The *R. shameli* ACE2 was used as the outgroup for rooting the tree. The mutations of each branch including nucleotide mutations (top) and corresponding amino acid mutations (bottom) are annotated on the branches and font color is different for non-synonymous (dark) and synonymous (light) changes. The substitutions that define northern/southern lineage are emphasized with coloured boxes. Tips are coloured based on the locations where the sequences were sampled at. Tips at parts of the tree with more than 5 identical nucleotide sequences sampled at the same location are collapsed and displayed as diamonds with the number of identical sequences written on the right of each shape. The tree is categorized into three parts in accordance with their phylogenetic clustering: northern lineage including 46 sequences and 9 genotypes, southern lineage comprising 82 sequences and 4 genotypes, and *R. affinis* basal clade with the remaining 54 sequences. The expanded tree is displayed in **Figure S2B**.

We reconstructed a phylogenetic tree of the 155 CYTB sequences derived from liver samples together with publicly available sequences of related *Rhinolophus* species to confirm species identification **(Figure S1)**. The resulting phylogeny clearly demonstrates high genetic similarity between our sequences, which cluster monophyletically with available *R. affinis* CYTB sequences. Moreover, the sequences form two distinct genetic clusters, consistent with the geography of the bats’ sampling locations. Samples collected from Bach Ma in the southern region predominantly fall within one distinct clade, whereas samples from Tam Dao and Lang Son in the northern region cluster separately into another clade **(Figure S1)**. Interestingly, samples from the Cat Ba Island exhibit a mixed pattern, being distributed across both clades, and 2 individuals from Bach Ma were clustered in the predominantly Tam Dao and Lang Son clade.

### Genetic characterization and polymorphism of ACE2 in *R. affinis*

We PCR-amplified and sequenced ACE2 transcripts obtained from organ samples of all 155 *R. affinis* individuals. After excluding false positives through molecular cloning, 159 ACE2 transcript sequences were isolated, indicating the presence of double haplotypes in four individuals, as detailed in **Tables S1 and S2**.

Building upon previous research^24^, we have considerably expanded the existing dataset of *R. affinis* ACE2 sequences. The 159 ACE2 coding sequences derived from 155 individuals presented here correspond to 30 unique nucleotide sequences and 15 unique amino acid genotypes. Additionally, we retrieved 23 complete *R. affinis* ACE2 coding sequences^24^, corresponding to 7 distinct amino acid genotypes. Five amino acid genotypes retrieved from GenBank matched genotypes identified in our dataset. Consequently, a total of 10 novel ACE2 genotypes were characterized from the 155 *R. affinis* individuals collected in Vietnam as part of this study. Following alignment and genotype assignment of all coding sequences, one genotype consistently found in publicly available sequences (GenBank accession QMQ39222) and in our dataset was designated as “V-Ra-01”, serving as a reference for naming subsequent genotypes, as indicated in **Table S2**. Furthermore, to comprehensively characterize ACE2 polymorphisms within *R. affinis*, we integrated two additional genotypes exclusively reported from prior studies conducted in China (also renamed from “RA-01” and “RA-02” to “C-Ra-01” and “C-Ra-02” in our study, respectively)^24^, ultimately resulting in a genotype dataset encompassing 17 unique ACE2 protein sequences.

We identified 9 protein sites varying within the available *R. affinis* ACE2 sequences. Most of the sites fall on the N-terminal region of the ACE2 protein and follow a consistent pattern in residue combinations. Interestingly, the residue distribution of site 603 (the most C-terminal variable site) was seemingly inconsistent to that of the other variable sites, implicating the possibility of genetic recombination. To further explore the residue distribution, we inferred a phylogenetic tree based on the full ACE2 coding sequences and mapped nucleotide substitutions onto the tree branches **(Figure S2A)**. Multiple recurrent nucleotide substitutions were observed on the codon corresponding to residue 603, supporting that the residue distribution on this amino acid site may have been influenced by recombination. Therefore, we conducted recombination analysis using the genetic algorithm for recombination detection (GARD)^33^ and identified a strongly supported breakpoint at nucleotide position 1685, indicating a recombination event during the evolutionary history of the *R. affinis* ACE2 gene **(Figure 2C)**. Based on this result, the aligned ACE2 coding sequences were partitioned into two segments at the detected breakpoint position. Each alignment partition was then used to infer phylogenies and branch-specific substitutions in order to remove the recombination signal **(Figures 1B, S2B and S2C)**.

**Figure 2.**
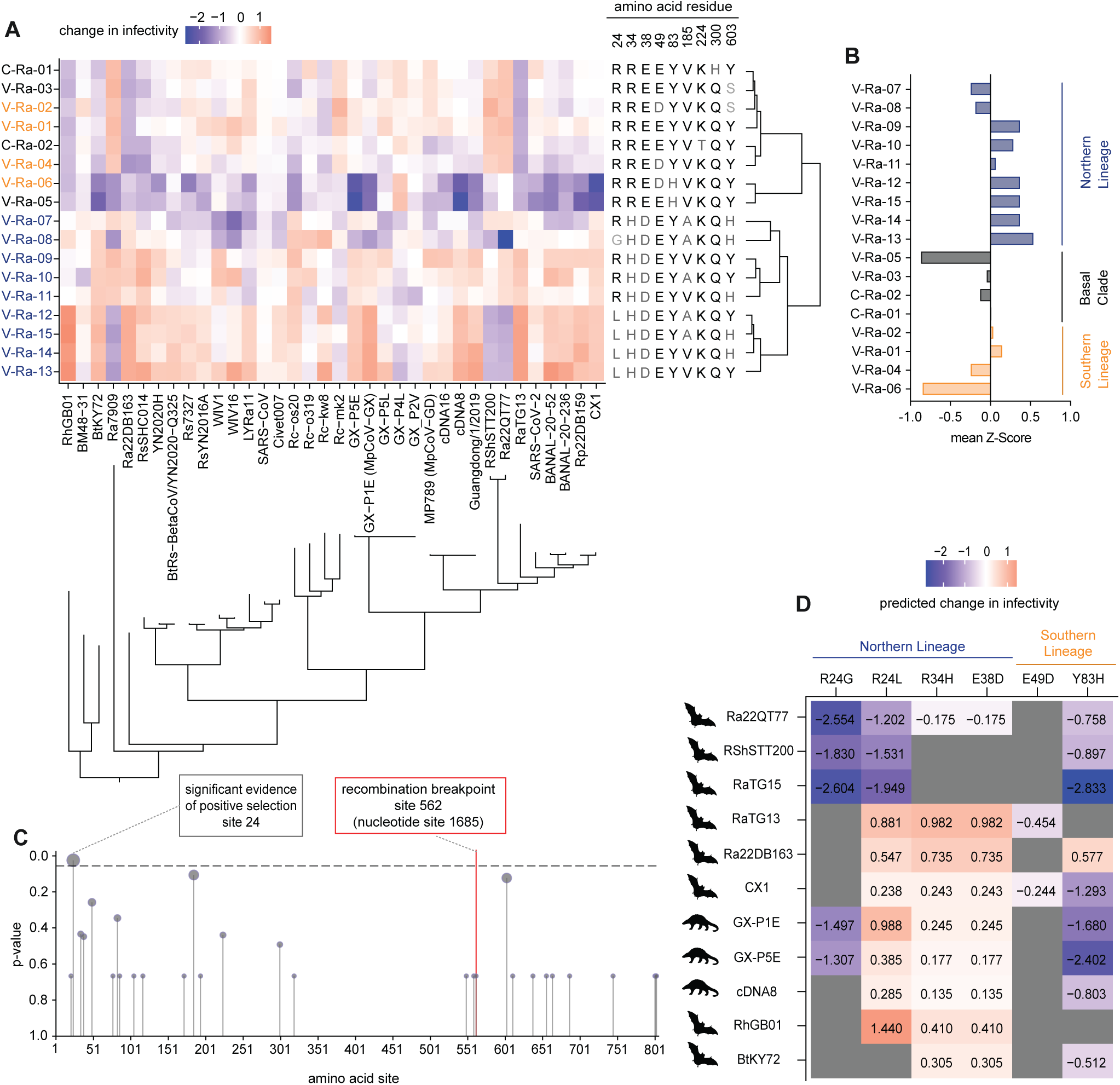
Genetic features and susceptibility profile of *R. affinis* ACE2. **(A)** Heatmap of infectivity profiles across HOS-ACE2/TMPRSS2 cell lines and pseudotyped sarbecovirus spikes. Rows show 17 HOS-ACE2/TMPRSS2 cell lines and columns show 36 sarbecovirus spikes. For each virus, infectivity values across the 17 cell lines were normalized and scaled to a z score, which is mapped to a diverging colormap. Higher z scores appear in red and indicate greater relative susceptibility of the ACE2 cell lines; lower z scores appear in blue and indicate reduced susceptibility. ACE2 genotypes are coloured based on their geographic lineages, consistent with **Figures 1B and 2B**. For each ACE2 genotype, amino acid residues at variable sites are annotated alongside the heatmap. The dendrogram on the right of the heatmap corresponds to the similarity between infectivity profiles. A maximum likelihood tree of the sarbecoviruses is displayed beneath the heatmap. **(B)** Mean value of z-score for each ACE2 genotype corresponding to the heatmap in plan A. The 17 ACE2 genotypes were classified into 3 lineages and plotted with different colors consistent with **Figures 1B and 2A**. **(C)** Lollipop plot of p-values corresponding to significance of evidence for positive selection determined by MEME. The horizontal line marks the significance threshold (*p* ≤ 0.05). The position corresponding to the recombination breakpoint determined by GARD is annotated on the plot. **(D)** Heatmap of regression coefficients quantifying the effect of 6 ACE2 amino acid changes (columns; across 17 ACE2 genotypes) on normalized infectivity for 36 sarbecoviruses (rows; annotated by host origin). Cell values are log-scaled regression coefficients: blue indicates decreased infectivity and red increased infectivity; non-significant associations (*p* ≥ 0.05) are shown in dark gray. Eleven of the 36 viruses exhibited at least one significant association (*p* < 0.05). See more details in Methods.

Since the non-recombinant partition representing the C-terminus of the ACE2 is much shorter and contains a lot fewer informative sites **(Figures 2C and S2C)**, only site 603 being variable on the protein level, we decided to focus our phylogenetic analysis on the partition representing the N-terminal region of the protein. As shown in **Figure 1B**, most sequences previously sampled from China, some sequences from Bach Ma and 1 from Cat Ba fall on the base of the *R. affinis* clade (basal clade, encoding for genotypes V-Ra-01, V-Ra-03, V-Ra-05, C-Ra-01 and C-Ra-02), while the rest of the genotypes broadly segregated into two distinct lineages corresponding to the geographic locations where the bats were sampled: The northern lineage (including all bats from Tam Dao and Lang Son, 2 from Cat Ba, and only 1 from Bach Ma) is defined by two non-synonymous substitutions g101a and g114c, leading to protein changes R34H and E38D; The southern lineage (comprising of most bats from Bach Ma and 1 from Cat Ba) is defined by the synonymous substitution c957g **(Figure 1B)**. Furthermore, this geographically structured topology closely aligns with that of the CYTB phylogeny **(Figure S1)**, underscoring the geographical separation of the sampled *R. affinis* populations. Interestingly, genotype V-Ra-01 occupies the most basal position in the phylogenetic tree and exhibits the widest geographic distribution, extending from Hubei Province, China, to southern Vietnam **(Figures 1B and S2B)**. Such similar geographic distribution pattern is also evident in four additional genotypes: V-Ra-04 and V-Ra-05 from southern Vietnam, and V-Ra-10 and V-Ra-11 from northern Vietnam, all of which have also been previously identified in Yunnan Province, China. Furthermore, the two ACE2 genotypes only sampled in China so far (C-Ra-01 and C-Ra-02) exhibit the closest genetic relatedness with genotype V-Ra-01.

Despite considerable diversity within ACE2 genotypes observed even within the same sampling location, the overall distribution pattern displayed clear geographical separation **(Figure S3A).** Without accounting for overlaps, we independently identified 7, 6, 8, and 3 genotypes from Bach Ma, Tam Dao, Lang Son, and Cat Ba, respectively, highlighting notable ACE2 polymorphism within *R. affinis*. Moreover, several genotypes were observed across multiple locations (e.g., genotype V-Ra-10), whereas others appeared exclusively within specific regions, such as genotype V-Ra-02 in Bach Ma, V-Ra-12 in Tam Dao, and V-Ra-14 in Lang Son **(Figure S3A)**. Notably, all genotypes from the Cat Ba Island **(Figure 1A)**, were shared among the other three mainland locations, without any unique genotype exclusive to the island **(Figure S3A and Table S2)**. This separation is consistent with the CYTB analysis results **(Figure S1),** suggesting potential gene flow between populations of *R. affinis*, despite the geographic isolation of Cat Ba Island.

In summary, we have categorized all 17 identified ACE2 genotypes into two distinct lineages according to their phylogenetic clustering: the northern lineage (comprising 9 genotypes) and the southern lineage (consisting of 4 genotypes). The remaining 4 genotypes (V-Ra-03, V-Ra-05, C-Ra-01, C-Ra-02) fall on the base of the *R. affinis* clade and are genetically closest to genotype V-Ra-01 sequences **(Figure 1B)**.

### Distinct sarbecovirus susceptibility between *R. affinis* ACE2 genotypes

To investigate whether the characterized ACE2 polymorphism has an effect on how each genotype interacts with ACE2-using sarbecoviruses, we constructed HOS - human TMPRSS2 cell lines over-expressing each of the 17 ACE2 genotypes (HOS-ACE2/TMPRSS2 cell lines; **Figure S3B**) and infected them with pseudoviruses carrying a panel of 36 sarbecovirus spike proteins. These include bat sarbecoviruses sampled in *R. affinis*, *R. shameli*, *R. pusillus*, *R. sinicus, R. marshalli, R. hipposideros, R. blasii, R. cornutus* and *R. malayanus* as well as pangolin sarbecoviruses, SARS-CoV-2, and SARS-CoV from humans and civets **(Table S3)**. We only selected viruses previously classified into sarbecovirus clades 1 and 3 rather than clade 2 since the latter are incapable of using ACE2 as their cellular receptor according to previous studies^29^. Notably, the resulting clustering of viral tropism also displayed clear geographical segregation, distinguishing the northern from the southern lineage (as well as basal genotypes V-Ra-03, V-Ra-05, C-Ra-01, C-Ra-02) **(Figures 2A and 2B)**.

Within the basal clade and southern lineage, genotypes harbouring the Y83H mutation (V-Ra-05 and V-Ra-06) exhibited reduced susceptibility to nearly all tested sarbecoviruses, suggesting a narrower infectivity profile compared to other *R. affinis* ACE2s. This observation aligns closely with previously reported findings^24^. Conversely, within the northern lineage, genotypes carrying a leucine at residue 24 (L24) appeared to have enhanced susceptibility toward multiple sarbecoviruses **(Figure 2A)**. This effect is clearly illustrated by comparing the mean z-scores for all 36 sarbecoviruses, as shown in **Figure 2B**, between the following genotype pairs: V-Ra-07 and V-Ra-15 (R24L) as well as V-Ra-08 and V-Ra-15 (G24L), both of which showed a marked increase in susceptibility when containing a leucine at site 24. Similarly, other genotype pairs only differing in their site 24 residue (R24L: V-Ra-09 and V-Ra-13, V-Ra-10 and V-Ra-12, V-Ra-11 and V-Ra-14) exhibited modest yet consistent increases in susceptibility **(Figure 2B)**. It is worth noting that the divergence at residue 24 occurred exclusively within the northern ACE2 lineage (defined by substitutions R34H, and E38D), independently to leucine (R24L) and to glycine (R24G) **(Figure 1B)**.

Most *R. affinis* ACE2 genotypes exhibited a broad spectrum of susceptibility toward the extensive panel of tested sarbecoviruses **(Figure 2A and Data S1),** however, some viruses showed distinct tropism profiles against the tested ACE2s, particularly those originally sampled from *R. affinis* individuals. We selected 5 representative sarbecoviruses (Ra22QT77, RaTG13, RaTG15/Ra7909, Ra22DB163 from *R. affinis* and RshSTT200 from *R. shameli*, **Table S3**) to further investigate these unique patterns. Analogous to the ACE2 lineages, these sarbecoviruses can be divided based on their sampling locations: RaTG13 and RaTG15 from Yunnan, China; Ra22DB163 from Dien Bien, Vietnam; Ra22QT77 from Quang Tri, Vietnam; and RShSTT200 from Stung Treng, Cambodia **(Figure 1A and Table S3)**. Remarkably, we observed evident geographic differences in viral tropism between these groups. The viruses sampled in Yunnan, China and Dien Bien, Vietnam (north) preferentially infected northern ACE2 lineage, whereas the viruses sampled in Quang Tri, Vietnam and Stung Treng, Cambodia (south) demonstrated a stronger affinity toward southern lineage ACE2s (and basal genotypes V-Ra-03, V-Ra-05, C-Ra-01, C-Ra-02) **(Figure 2A and Data S1)**.

Additionally, ACE2 genotypes found in China exhibited infectivity profiles remarkably similar to those of ACE2s sampled in southern Vietnam **(Figure 2A and Data S1)**. Notably, this phenotypic consistency closely aligns with our phylogenetic tree **(Figure 1B)**, suggesting that southern Vietnam contains the most evolutionarily basal genotypes of *R. affinis* ACE2, whereas northern Vietnam ACE2s seem to be geographically, genetically, and phenotypically separated from the others **(Figures 1A, 1B, and 2A)**.

### Quantitative evaluation of ACE2 substitutions’ effect on virus susceptibility

Given the phenotypic variation between *R. affinis* ACE2s **(Figure 2A)**, we explored whether any sites show evidence of positive selection. Among the 32 sites that are variable on the nucleotide level across our ACE2 dataset contains 21 synonymous substitutions, and 11 non-synonymous substitutions **(Figure 1B)** which subsequently lead to variation on the protein level. We tested for site-specific diversifying selection within the *R. affinis* ACE2 evolution using the mixed effects model of evolution (MEME) method^34^. This was done separately for each non-recombinant partition defined above **(Figure S2)** to avoid the influence of recombination. The only position in the protein that showed significant evidence for diversifying positive selection in either partition was site 24 (*p* = 0.025) **(Figure 2C)**. As shown in **Figure 1B**, this signal of positive selection is supported by two independent non-synonymous substitutions on site 24 within the *R. affinis* ACE2 phylogeny (R24G resulting from a single c70g nucleotide substitution, and R24L resulting from two nucleotide substitutions c70t and g71t), both occurring within the northern ACE2 lineage.

To quantitatively evaluate the phenotypic impact associated with each varying site, we performed a linear regression analysis, quantifying the effect each residue has on the infectivity of each tested sarbecovirus. As illustrated in **Figure 2D**, our analysis showed that the infectivity profile of 11 out of the 36 sarbecoviruses examined was significantly affected by the ACE2 polymorphism. Notably, these 11 sarbecoviruses originated from bats (*R. affinis*, *R. shameli*, *R. marshalli, R. pusillus* and *R. hipposideros*) as well as pangolins (*Manis javanica*) **(Table S3)**.

Furthermore, we observed that Ra22QT77 and RshSTT200, previously identified in southern Vietnam and Cambodia **(Figure 1A)** and exhibiting a close genetic relationship to one another **(Figure 2A)**, demonstrated similar tropism associated with specific ACE2 residues, particularly at site 24. Both viruses displayed reduced infectivity toward mutations occurring in the northern ACE2 lineage (R24L decreased by 1.2-1.6 log and R24G decreased by 1.8-2.6 log, **Figure 2D**), which aligns well with the geography of their sampling locations. RaTG15 was another virus that showed a highly similar infectivity pattern to the aforementioned two sarbecoviruses. However, RaTG15 differs substantially in its phylogenetic and geographic placement compared to Ra22QT77 and RshSTT200 **(Figures 2A and 1A)**. This divergence suggests that RaTG15 likely evolved this ACE2 tropism independently, acquiring additional mutations to preferentially adapt to ACE2 genotypes prevalent in the southern lineage.

On the other hand, RaTG13 and Ra22DB163, sampled in Yunnan, China and Dien Bien, Vietnam **(Figure 1A)**, demonstrated enhanced infectivity associated with mutations specific to the northern ACE2 lineage (R24L increased by 0.5-0.9 log and R34H/E38D increased by 0.7-1.0 log, **Figure 2D**), consistent with their sampling locations. The 3 sarbecoviruses sampled in pangolins (GX-P1E, GX-P5E, and cDNA8) also exhibited significantly enhanced infectivity when the amino acid at site 24 was substituted from arginine to leucine (R24L increased by 0.2-1.0 log; **Figure 2D**). Conversely, GX-P1E and GX-P5E showed significantly reduced infectivity following a substitution at the same site from arginine to glycine (R24G decreased by 1.3-1.5 log; **Figure 2D**).

### Specialization of Ra22QT77 to distinct *R. affinis* ACE2 genotypes

RshSTT200 and Ra22QT77 are of particular interest, being the only known clade 1b viruses incapable of utilizing human ACE2 as a receptor for infection^29^. Compared to the SARS-CoV-2 spike protein, the spike proteins of Ra22QT77 and RshSTT200 both exhibit two deletions at residues 426-427 and 430-431, and four key amino acid substitutions (T328R, P468F, D469N, and Y478G), all of which are located within the ACE2-binding interface^28^. Furthermore, reverting these substitutions into the RBD of RshSTT200 restores its ability to use human ACE2 for viral entry^29,35^.

Building upon these findings, we hypothesized that the observed sequence differences (i.e., the two deletions and four amino acid substitutions) between SARS-CoV-2 and Ra22QT77 (as well as RshSTT200) may have arisen because of adaptive evolution in response to selective pressures exerted by different *R. affinis* ACE2 genotypes. Given that Ra22QT77 was sampled in *R. affinis* from a site that is geographically close to our Bach Ma site, it is reasonable to propose that these genetic features of the virus were shaped primarily by interactions with host ACE2 receptors during its evolution and circulation. To test this hypothesis, we generated a series of pseudoviruses with the spike proteins of Ra22QT77 derivatives, including variants with restored deletions (i.e., corresponding insertions from the perspective of Ra22QT77), four individual amino acid substitutions, and combinations thereof. Infectivity assays were subsequently performed using these Ra22QT77 mutants to infect 17 HOS-ACE2/TMPRSS2 cell lines accordingly.

As shown in **Figure 3A**, the Ra22QT77 derivative harboring both restored deletions and all four substitutions (Ra22QT77 ins 4sub) exhibited reduced infectivity toward southern lineage ACE2s (as well as basal clade genotypes V-Ra-03, V-Ra-05, C-Ra-01, C-Ra-02), while its infectivity towards northern lineage ACE2s was only slightly reduced relative to the wild-type Ra22QT77 spike. Furthermore, the mutant containing the four amino acid substitutions (Ra22QT77 4sub) showed generally higher infectivity across nearly all ACE2 genotypes compared to both the wild-type Ra22QT77 spike and the Ra22QT77 ins 4sub mutant, having a more global effect on *R. affinis* ACE2 usage.

**Figure 3.**
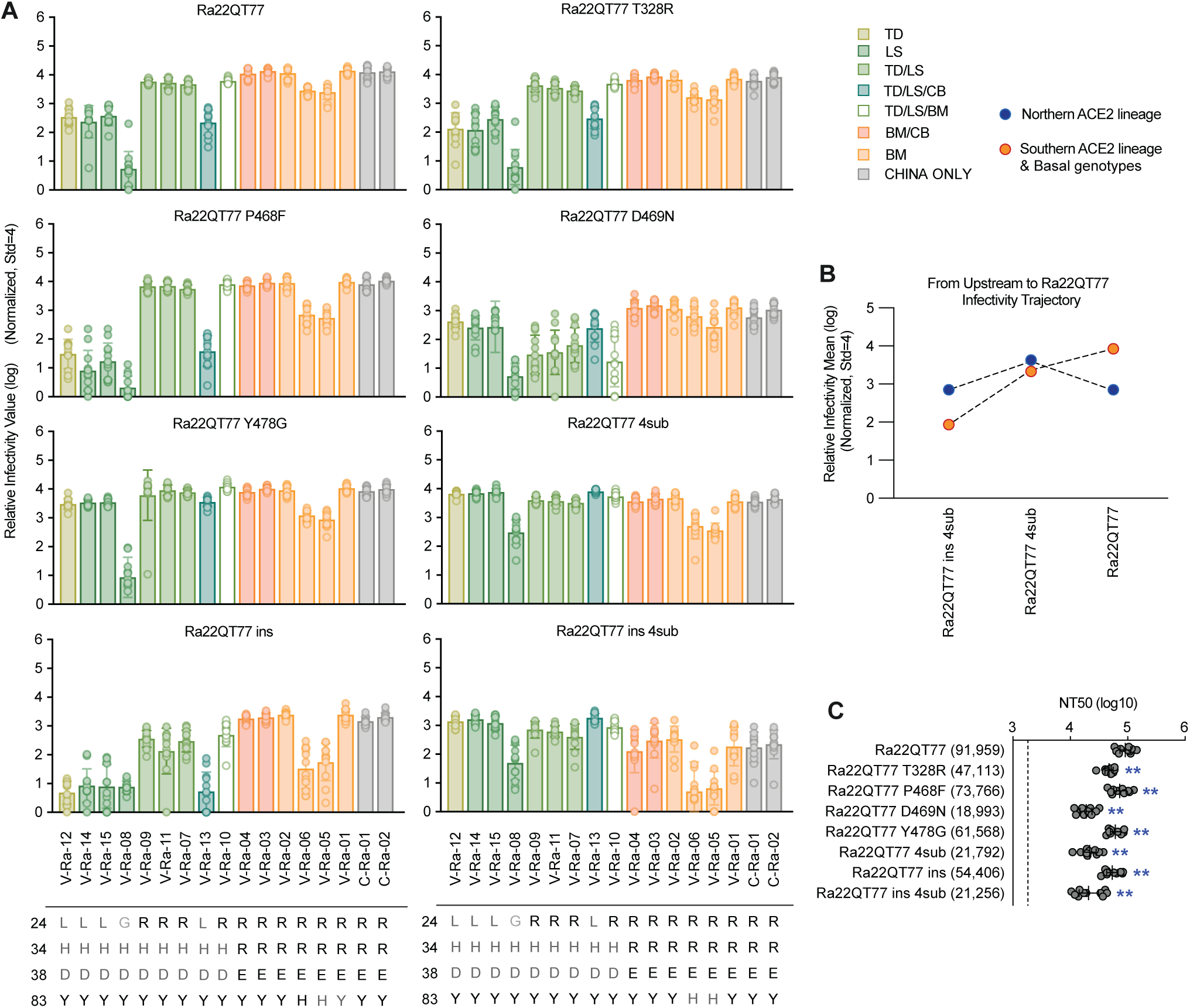
Specialization of Ra22QT77 derivatives to distinct *R. affinis* ACE2 genotypes. **(A)** Infectivity plots of 17 *R. affinis* ACE2 genotypes infected with Ra22QT77 and derivatives. The infectivity data of each panel is normalized by an internal infection control (HOS-TMPRSS2 cell line with human ACE2 receptor infects with SARS-CoV-2 spike) and displayed as log value. Each bar plot includes 12 replicates from 3 trials. The genotypes are ordered and categorized into 8 location-associated groups. TD, LS, CB, BM are abbreviations of Tam Dao, Lang Son, Cat Ba, and Bach Ma, respectively. The amino acid differences on residues 24, 34, 38, and 83 were attached beneath the infectivity plots. **(B)** Line plot summarizing normalized infectivity **(Figure 3A)** across 17 ACE2 genotypes classified into two groups, ordered along a hypothesized path from a proximal ancestror to Ra22QT77. Group means are plotted at each position. Three focal derivatives are shown: Ra22QT77 ins 4sub (putative ancestral RBD), Ra22QT77 4sub (intermediate), and Ra22QT77 (derived). **(C)** Neutralization assay of Ra22QT77 derivatives. Neutralization assays were performed with pseudoviruses harboring the spike proteins of Ra22QT77 derivatives against mouse sera elicited by 3-dose 10 ug Ra22QT77 spike mRNA vaccination (n = 10). The NT50 value of all Ra22QT77 variants are performed Wilcoxon test with Ra22QT77 (two-tailed test) and the results are all with significant difference. *: *p* < 0.05, **: *p* < 0.01, ***: *p* < 0.001.

By comparing the spikes differing only in the presence of the two deletions (i.e., Ra22QT77 ins 4sub and Ra22QT77 4sub; Ra22QT77 ins and Ra22QT77; **Figure 3A**), it is clear that the deletions contribute to an expanded viral tropism, enabling more efficient utilization of different *R. affinis* ACE2 receptors from various bat populations **(Figure 3B)**. Conversely, comparing spikes differing only in the four amino acids (i.e., Ra22QT77 ins 4sub and Ra22QT77 ins; Ra22QT77 4sub and Ra22QT77; **Figure 3A**) shows that these substitutions enhance the virus’s ability to use southern lineage ACE2s and basal clade genotypes **(Figure 3B)**.

Despite their clear effect on ACE2 usage, it remains uncertain whether these spike changes were driven exclusively by receptor-mediated selection, and in which order the deletion and substitution events took place. If the proximal ancestor of these viruses acquired these changes as a result of adaptation to a new *Rhinolophus* host, then adapting to more efficiently using the host’s receptor is expected to happen first. Once circulation continues and immunity builds up in the population, adaptive changes in the virus RBD are more likely to occur as a response to immune pressure. To test this hypothesis, we immunized mice with 3 doses of a Ra22QT77 spike-encoding mRNA vaccine and performed neutralization assays using the collected sera against each Ra22QT77 mutant spike described above. Results indicated significantly lower NT50 values for all mutants compared to the wild-type Ra22QT77 spike (*p* = 0.002-0.006), suggesting enhanced resistance to neutralization by anti-Ra22QT77 sera **(Figure 3C)**. However, spikes differing only by the two deletions only showed a very subtle effect in neutralization titer (Ra22QT77 ins 4sub and Ra22QT77 4sub decreased by 0.01 log of NT50; Ra22QT77 ins and Ra22QT77 decreased by 0.23 log of NT50). In contrast, spikes with all four amino acid substitutions or particularly substitution D469N had a much more pronounced effect on neutralization titers (Ra22QT77 4sub and Ra22QT77 decreased by 0.63 log of NT50; Ra22QT77 D469N and Ra22QT77 decreased by 0.69 log of NT50) **(Figure 3C)**. Collectively, these findings suggest that the two deletions likely happened first as a result of receptor-specific pressure, and the four substitutions followed as a result of immune pressure.

### Molecular interactions between *R. affinis* ACE2 and the Ra22QT77 RBD

To further elucidate the molecular interactions underlying the Ra22QT77 RBD, we resolved the complex structure of the RBD bound to the virus’s natural host receptor, *R. affinis* ACE2 (V-Ra-10), using cryo-EM **(Figures 4A and S4 and Data S2)**. Structural analysis indicated that R24 in *R. affinis* ACE2 directly contacts site D469 in the Ra22QT77 spike via two non-hydrogen-bond interactions. Given the positive charge of the arginine and the negative charge of aspartic acid, this interaction likely forms salt bridges, providing a substantial electrostatic contribution to binding affinity **(Figure 4B)**. This finding aligns with our infectivity assays, which showed markedly decreased susceptibility of ACE2s harboring R24L or R24G substitutions **(Figure 2D)**. Hence, substituting arginine with leucine at site 24 eliminates electrostatic interactions and hydrogen-bonding capabilities, retaining only hydrophobic compensation, thereby weakening binding. The R24G substitution further reduces affinity by entirely removing the side chain, abolishing all favorable binding interactions.

**Figure 4.**
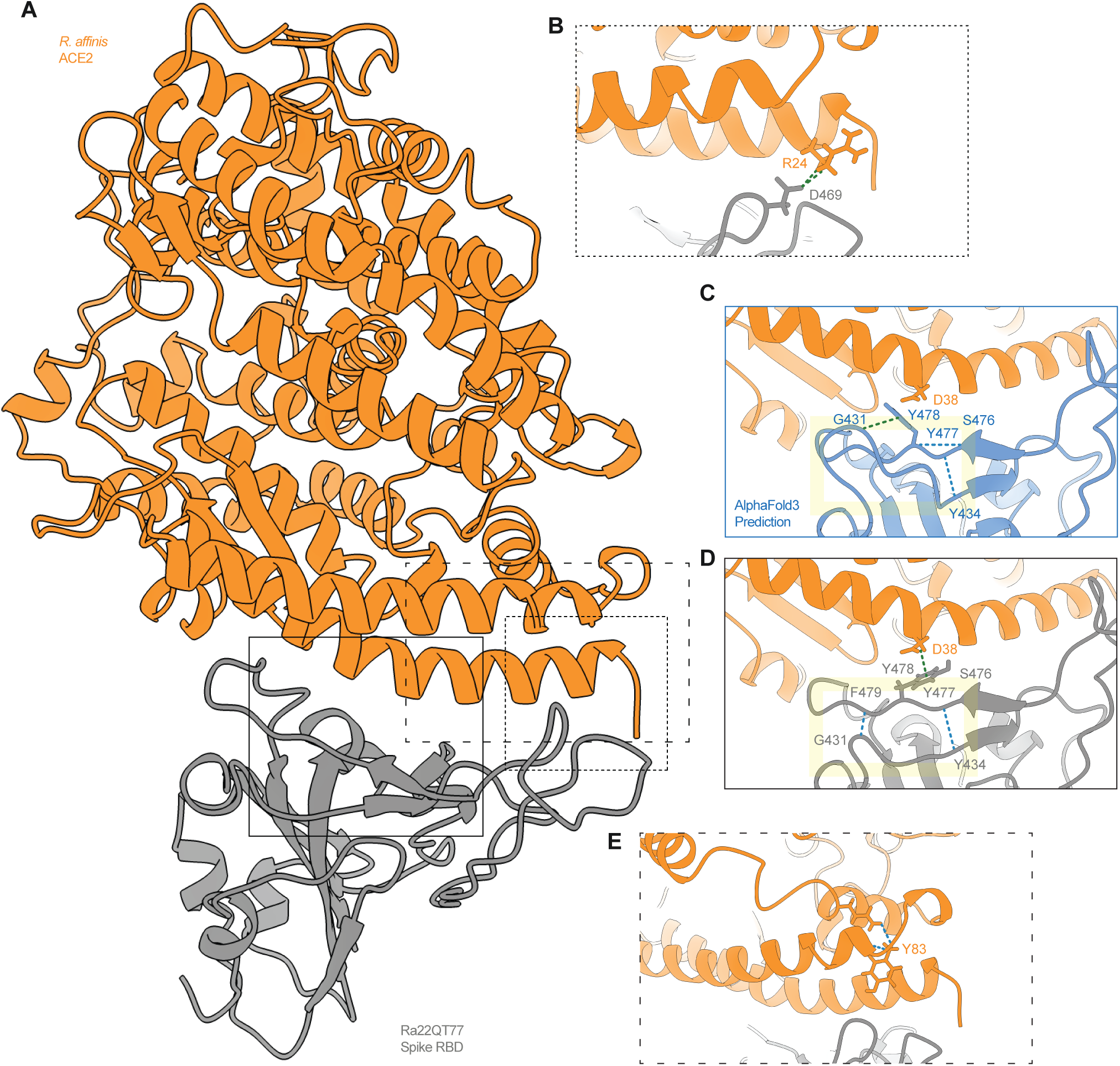
Molecular interactions of Ra22QT77 spike RBD and *R. affinis* ACE2 with cryo-EM and AlphaFold 3 prediction. **(A)** The cryo-EM structure of the complex of Ra22QT77 spike RBD and *R. affinis* ACE2. The 3-dimension structures of Ra22QT77 spike RBD and *R. affinis* ACE2 (genotype: V-Ra-10) are plotted as grey and orange, respectively. Areas of interest are annotated in close-up views shown in **Figure 4B, 4D, and 4E**. The outline style of each close-up window corresponds to the outline style of the respective panel B, D, and E. The residue numbering is based on Ra22QT77 spike RBD sequence. **(B)** The non-hydrogen-bond interactions between D469 from Ra22QT77 spike RBD and R24 from *R. affinis* ACE2 are shown as green dashed lines and inferred as salt bridge interactions. **(C)** AlphaFold3 predicted structure of the Ra22QT77 ins RBD. Residue numbering is consistent with that of the wild-type Ra22QT77 RBD (panel D). The hydrogen bonds between the following groups are shown as blue dashed lines: S476 and Y478, and Y477 and Y434. The non-hydrogen-bond interactions between Y478 and G431 are merged and shown as green dashed line and inferred as van de Waals contacts. The local loop structure with the 4 amino acid deletion restored is highlighted in a yellow box. **(D)** The hydrogen bonds between Y477 and Y434, and F479 and G431 are shown as blue dashed lines. The non-hydrogen-bond interactions between Y478 from Ra22QT77 spike RBD and D38 from *R. affinis* ACE2 are merged and shown as green dashed line and inferred as van de Waals contacts. The local loop structure is highlighted in a yellow box. **(E)** The hydrogen bonds among Y83 and other spatial-adjacent residues from *R. affinis* ACE2 are shown as blue dashed lines.

Regarding the two deletions present on the RBD of Ra22QT77 and RshSTT200, we compared our cryo-EM structure **(Figure 4A)** to the AlphaFold3 predicted co-structure where the deletions were restored to investigate their potential molecular effect **(Figures 4C and 4D)**. Restoring K426, V427, N430 and Y431, showed that residue Y434 forms a hydrogen bond with Y477, and S476 forms another hydrogen bond with Y478 within the spike protein, while G431 engages in van der Waals contacts with Y478 **(Figure 4C)**. The residue numbering here corresponds to sites without restoration to keep consistent with cryo-EM structure **(Figure 4D)**. Deletion of the four residues in the receptor binding loop reorganizes this interaction network, The cryo-EM structure shows that Y434 still maintains its hydrogen bond with Y477, yet G431 shifts its spatial position which loses the contacts with Y478 and forms an additional hydrogen bond with F479 **(Figure 4D)**. These structural rearrangements effectively stabilize the spatial position of Y478 and increase the residue’s accessibility, thereby enhancing its ability to recognize and bind to residue 38 of *R. affinis* ACE2. Notably, Y478 is able to engage in van der Waals contacts with residue 38 on the ACE2 regardless of whether this position is occupied by aspartic acid (D38) or glutamic acid (E38). This could explain why the deletions enhance the capacity of Ra22QT77 spike to recognize and bind to different ACE2 lineages **(Figures 3A and 3B).**

On the other hand, our cryo-EM data explains why the Y83H mutation in *R. affinis* ACE2 broadly reduces infectivity of multiple sarbecoviruses **(Figures 2A, 2D)**. In the *R. affinis* ACE2, Y83 participates extensively in an intramolecular hydrogen-bond network; its phenolic ring forms aromatic stacking interactions with adjacent residues, and its hydroxyl group functions both as a hydrogen-bond donor and acceptor, crucial for maintaining local structural integrity. Notably, Y83 does not directly contact the Ra22QT77 spike protein **(Figure 4E).** Upon mutation to histidine (Y83H), the smaller, more polar imidazole group with partial positive charge at physiological pH replaces the phenolic ring, diminishing aromatic stacking interactions. This substitution likely disrupts intramolecular hydrogen bonds, alters local packing, and induces subtle conformational changes in ACE2, thereby destabilizing the structure. Such structural perturbations decrease ACE2 binding affinity, consequently reducing viral attachment efficiency across different sarbecovirus spikes.

## Discussion

The ACE2 protein shows pronounced intraspecific polymorphism in horseshoe bats, which has been reported in *Hipposideros armiger*, *R. sinicus*, *R. ferrumequinum* and *R. affinis*^24–26^. However, it remained unclear what the drivers of this genetic diversity are, as well as how this receptor diversity affects how different horseshoe bat populations interact with circulating sarbecoviruses. To answer these questions, we collected 155 *Rhinolophus affinis* samples in Bach Ma, Tam Dao, Lang Son and Cat Ba, Vietnam during 2021 and 2022 and identified 15 genotypes of *R. affinis* ACE2. We show that ACE2 polymorphism is geographically separated and associated with distinct sarbecovirus entry phenotypes by phylogenetic analysis and infectivity assays. Focusing on the interaction between *R. affinis* ACE2 and representative sarbecovirus Ra22QT77 (sampled in the same host species), we identify mutations controlling the infectivity phenotype on both the receptor and the virus spike protein. Hence, we present evidence of both host receptor polymorphism shaping virus evolution but also, perhaps more interestingly, ancestral sarbecoviruses likely having shaped the intra-species evolution of horseshoe bat ACE2 proteins.

Our ACE2 data reveal a clear north-south divide within the sampled populations of *R. affinis*. Individuals sampled in northern Vietnam form a distinct clade based on both their mitochondrial CYTB and ACE2 sequences, while exhibiting distinct sarbecovirus infectivity profiles **(Figures 1B, 2A, and S1)**. This northern ACE2 lineage is defined by two non-synonymous substitutions, R34H and R38D **(Figure 1B)**, both of which significantly alter the ACE2s’ sarbecovirus infectivity profiles **(Figure 2D)**. Within the northern lineage we document two independent substitutions at site 24 (R24L and R24G; **Figure 1B**), both of which lead to large changes in susceptibility to multiple sarbecoviruses (both increases and decreases; **Figure 2D**). Independent of our infectivity results, we detect significant evidence of positive diversifying selection on site 24 within the evolution of the *R. affinis* ACE2 (**Figure 2C**). Taken together, these findings suggest that infection by an ancient sarbecovirus likely shaped the genetic polymorphism of ACE2 within this horseshoe bat species, represented by the geographically separated northern Vietnam population we have sampled as part of this study. It is also worth noting that south Vietnam (Bach Ma) ACE2s include genotypes from the basal clade of the phylogeny (**Figure 1B**) which are both genetically and phenotypically more similar to the ACE2 genotypes sampled in China (**Figure 2B**). This further implies that the proposed ancestral sarbecovirus pressure is unique to the northern Vietnam *R. affinis* population. Even though we cannot reconstruct this putative sarbecovirus, we hypothesize that its receptor tropism resembled contemporary viruses such as Ra22QT77, RshSTT200, or RaTG15, whose infectivity can be reduced by the observed site 24 substitutions (and partly by the site 34 and 38 substitutions; **Figure 2D**). To our knowledge, this is the first direct evidence of sarbecovirus infection acting as a selective pressure on the receptor of the viruses’ natural reservoir host, potentially shifting the paradigm that these viruses cause little to no disease in the horseshoe bats they circulate in^36,37^.

Among the diverse panel of sarbecoviruses tested, several displayed strikingly similar infectivity patterns. Despite considerable sequence divergence between their RBDs, sarbecoviruses RaTG13 and Ra22DB163, demonstrated nearly identical entry efficiencies across the *R. affinis* ACE2s, preferentially binding to northern lineage genotypes **(Figures 2A and 2D)**. The RaTG13 and Ra22DB163 RBDs cluster into different parts of the sarbecovirus tree **(Figure 2A)**, estimated to have diverged thousands of years ago^38^. Nonetheless, each virus appears to have independently acquired substitutions in their RBDs that allows them to converge into near identical *R. affinis* ACE2 infectivity phenotypes **(Figure 2D)**. Similarly, Ra22QT77, RshSTT200 and RaTG15 show a very similar infectivity phenotype, in this case having much weaker binding to northern lineage ACE2s (**Figures 2A and 2D)**. Such convergence in RBD adaptations is broadly analogous to what has been observed for SARS-CoV and SARS-CoV-2^39^. It is evident that sarbecoviruses likely continually alter their infectivity profiles, depending, at least partly, on the bat population they infect^25,40,41^. This diversity in spike infectivity phenotypes and flexibility in switching between them showcases the importance of continuous virus surveillance paired with monitoring the receptor polymorphism of the host populations that the viruses are sampled in.

Using Ra22QT77 as an example, we were able to characterise the likely stepwise evolution of within host receptor specificity for this virus **(Figure 3B)**. This sub-group of virus RBDs have a characteristic four amino acid deletion along with a number of substitutions, that took place on the same branch of the RBD phylogeny and have only been sampled in Cambodia^28,42^ and Vietnam^38^. In our previous work, focusing on the use of human ACE2 by these viruses^29^ we find that reverting substitutions N469D and G478Y allows Ra22QT77 to infect hACE2, but the presence or absence of the four amino acid deletion has no effect on hACE2 usage. In this study we examine the effects of these substitutions on the usage of ACE2 genotypes within *R. affinis* bats in which these viruses naturally circulate. Interestingly, we find that the four amino acid deletion allows the Ra22QT77 spike to utilize a more diverse set of *R. affinis* ACE2 receptors (**Figures 3A and 3B**). Using cryo-EM and protein structure predictions we show that this deletion significantly alters the conformation of the elongated RBD loop, stabilizing the critical amino acid residue Y478, thereby enhancing its accessibility to the ACE2 receptor-binding interface (residues D/E38) **(Figures 4C and 4D)**. On the other hand, the four key RBD substitutions seem to decrease the binding of Ra22QT77 spike to primarily the northern lineage *R. affinis* ACE2s (**Figure 3B**). Instead, substitution D469N appears to have a pronounced effect on evading adaptive immunity (**Figure 3C**), whereas the other three mutations may compensate for the selective changes in the rest of the RBD **(Figures 3A and 3C)**. The geographic proximity of the sampling locations of viruses with these mutations (e.g., Ra22QT77 and RshSTT200) to our southern Vietnam Bach Ma site **(Figure 1A)** and apparent specific circulation in this area suggest a scenario wherein the mutations arose as a result of receptor usage specialisation to southern bat populations, followed by immune evasion once these viruses had spread through the new host population. This is further supported by the fact that the four tested substitutions moderately increase ACE2 binding to southern *R. affinis* ACE2s but reduce binding to northern ACE2s (where the viruses do not circulate) (**Figure 3B**).

Together, our findings provide important insights into the evolutionary dynamics shaping sarbecovirus diversity and host-virus interactions. Sarbecovirus spike proteins are fast-evolving and key sites in the RBD controlling receptor-binding constantly and frequently change throughout the viruses’ evolution^22,25^. Herein, we show that substantial polymorphism in the ACE2 receptor within the *R. affinis* species controls the spike evolution of sarbecoviruses circulating in these bats. Vietnam and other countries in Southeast Asia host the highest species richness of *Rhinolophus* bats, interacting with and transmitting viruses between each other^43,44^. Although, to date, little research has focused on bat ACE2 within-species diversity, we expect that similar levels of polymorphism will be present in other *Rhinolophus* species infected by the same viruses, collectively shaping sarbecovirus spike evolution. Just as ACE2 polymorphism affects how spike proteins evolve, our findings support that there is a bidirectional relationship between the two proteins’ evolution, where *R. affinis* ACE2’s evolution has also been shaped by selection from sarbecovirus infection. This is not a classical example of virus-host co-evolution, since the timescale of ACE2 evolution is much slower than that of the RBDs. However, we do observe phenotypic convergence in the spike proteins’ infectivity profiles, indicating continuous and complex interactions between the viruses and the bat receptors. Collectively, these findings stress the necessity of ongoing surveillance of both viral evolution and ACE2 diversity within reservoir host populations to predict potential zoonotic spillover events and better understand the processes underlying sarbecovirus circulation, emergence, and adaptation.

## Lead contact

Requests for further information, reagents, or resources may be directed to, and will be fulfilled by, the lead contact, Kei Sato.

## Data and code availability

Sequence data, metadata, phylogenies, raw experimental data, and code for the presented analysis are available in the following GitHub repository https://github.com/TheSatoLab/Raffinis_ACE2.

All ACE2 sequences will become available on DDBJ/ENA/GenBank upon paper acceptance.

The cryo-EM co-structure of the Ra22QT77 RBD bound to *R. affinis* ACE2 will become available on PDB upon paper acceptance under the accession 9X23 (EMDB-66471).

## Acknowledgements

We would like to express our gratitude to all the members of Division of Systems Virology, The Institute of Medical Science, The University of Tokyo. We thank Luo Chen, Dr. Alfredo Hinay Jr., and Dr. Jumpei Ito (The University of Tokyo) for providing insights on this work.

This study was supported in part by AMED ASPIRE program (25jf0126002 to Kei Sato); AMED SCARDA Japan Initiative for World-leading Vaccine Research and Development Centers “UTOPIA” (JP223fa627001 to Jumpei Ito, Spyros Lytras, and Kei Sato); AMED SCARDA Program on R&D of new generation vaccine including new modality application (253fa727002 to Kei Sato); AMED Research Program on Emerging and Re-emerging Infectious Diseases (24fk0108907, 25fk0108690 to Kei Sato); AMED Japan Program for Infectious Diseases Research and Infrastructure (Collaborative Research via Overseas Research Centers) (25wm0225041 to Kei Sato); JST PRESTO (JPMJPR22R1 to Jumpei Ito); JSPS KAKENHI fund for the Promotion of Joint International Research (International Leading Research) (JP23K20041 to Kei Sato); JSPS KAKENHI grant-in-aid for Scientific Research A (JP24H00607 to Kei Sato); JSPS research fellow DC1 (23KJ0710 to Yusuke Kosugi); JSPS research fellow DC2 (24KJ0628 to Shigeru Fujita); Mitsubishi UFJ Financial Group, Inc. Vaccine Development grant to Jumpei Ito and Kei Sato; the Platform Project for Supporting Drug Discovery and Life Science Research (Basis for Supporting Innovative Drug Discovery and Life Science Research [BINDS]) from AMED (JP24ama121002 (3272 to [Osamu Nureki]) and JP24ama121012 (S02820001 and S02820002 to Kei Sato).

## Author contributions

TT, NTT, LTQM, NTL, FH, and VDT conducted animal sampling and dissections;

WL, and SF performed extraction of RNA from tissues;

WL and DA conducted PCR amplification of ACE2 sequences;

WL and DA performed molecular cloning and stable cell line production;

WL and SF conducted PCR amplification of CYTB sequences;

WL, JET, and SL performed phylogenetic analysis;

WL performed selection and recombination analyses;

YK provided plasmids for pseudovirus assay;

WL constructed pseudoviruses and performed all infectivity assays;

XN prepared vaccinated murine sera;

MSY performed neutralisation experiments;

SS, HSO, KM, KS, and HA performed cryo-EM experiments and analysis;

WS, YC, FH, VDT, ON, SL, and KS supervised the work;

WL, SL, and KS conceptualised the study;

WL, and SL wrote the original manuscript;

All authors reviewed and proofread the manuscript.

## Declaration of interest

Spyros Lytras has previously received consulting fees from EcoHealth Alliance. Kei Sato has consulting fees from Moderna Japan Co., Ltd. and Takeda Pharmaceutical Co. Ltd., and honoraria for lectures from Moderna Japan Co., Ltd., Shionogi & Co., Ltd and AstraZeneca.

## Methods

### Ethics statement

The bat sampling conducted in Vietnam as part of this study has been approved by the ethics committee of the Nagasaki University Institute of Tropical Medicine (NEKKEN) (approval no. 200619239) and the National Institute of Hygiene and Epidemiology (NIHE) of Vietnam.

All mouse immunization experimental procedures were approved by the Animal Welfare Ethics Committee of Sinovac Biotech (protocol 202402-001).

### Sample collection and processing

Bats were collected using harp trap sampling and direct capture in caves. Chloroform was used for deep anaesthesia before euthanizing the animals. After anaesthesia, blood samples were drawn from heart, and the bodies were stored in ice separately and moved to the BSL2/3 facility in NIHE. Samples of brain, lung, heart, liver, spleen, intestinal tract, and kidney were taken from each body in the biosafety cabinet of the BSL2/3 facility, and all of the samples were stored separately at −80 C.

### Cell culture

LentiX-293T (a human embryonic kidney cell line; ATCC, Takara, Cat# 632180) and HOS cells (a human osteosarcoma cell line; ATCC CRL-1543) stably expressing human ACE2 and TMPRSS2^45,46^ were maintained in Dulbecco’s modified Eagle’s medium (high glucose) (Sigma-Aldrich, Cat# 6429-500ML) containing 10% fetal bovine serum and 1% penicillin-streptomycin (Sigma-Aldrich, Cat# P4333-100ML). The HOS-TMPRSS2 cells that stably express *R. affinis* ACE2 were generated as described below (see ‘‘**Generation of HOS-TMPRSS2 cells stably expressing R. affinis ACE2 proteins**’’ section) and maintained in Dulbecco’s modified Eagle’s medium (high glucose) (Sigma-Aldrich, Cat# 6429-500ML) containing 10% fetal bovine serum and 1% penicillin-streptomycin (Sigma-Aldrich, Cat# P4333-100ML), zeocin (50 μg/mL; InvivoGen, Cat# ant-zn-1) and G418 (400 μg/mL; Nacalai Tesque, Cat# G8168-10ML). Cell lines were tested for mycoplasma contamination by CycleavePCR Mycoplasma Detection Kit (Takara Bio, Cat# CY232).

### Nucleotide and amino acid sequences data collection

The spike sequences of sarbecoviruses used in this study were derived from our previous work^29^, with the exception of Ra22DB163. The Ra22DB163 spike sequences were retrieved from NCBI GenBank (Genbank ID: OR233325.1) and codon-optimized (Thermo Fisher Scientific). Also, the nucleotide sequences of all available *R. affinis* ACE2s were collected from NCBI GenBank (downloaded on 24^th^ of October 2024). The metadata of ACE2 sequences and information of the sarbecoviruses used in this study are summarized in **Tables S2 and S3**.

### ACE2 purification and amplification from bat organs

For each specimen, 50% of the total organ homogenate was allocated for RNA extraction. Samples were thawed on ice and homogenized prior to processing. 10 out of 159 ACE2 sequences were derived from lung tissues, 7 from intestinal tissues, and the remaining 142 from kidney tissues. Total RNA was extracted using the QIAamp RNA Blood Mini Kit (Qiagen, Cat# 52304) according to the manufacturer’s instructions. RNA concentration and purity were assessed with a NanoDrop spectrophotometer (Thermo Fisher Scientific). A portion of the purified RNA was reverse transcribed to cDNA using SuperScript III Reverse Transcriptase (Thermo Fisher, Cat# 18080085) in the presence of RNaseOUT Recombinant RNase Inhibitor (Thermo Fisher, Cat# EO038SKB011), following the supplier’s protocols and the specific primers designed. The resulting cDNA served as the template for three sequential rounds of nested PCR performed with PrimeSTAR Max DNA Polymerase (Takara Bio, Cat# R045A). For each round, primer pairs (synthesized by Fasmac, Atsugi, Japan) and cycling conditions were newly designed and empirically optimized. Amplicons were resolved by agarose gel electrophoresis using SeaKem GTG Agarose (Lonza, Cat# 50070). Target bands (*R. affinis* ACE2 DNA) were excised, and DNA was recovered and purified with a gel extraction kit (Qiagen, Cat# 28704) and tested with NanoDrop spectrophotometer (Thermo Fisher Scientific).

The primers used for this section are as follows:

ACE2_cDNA_syn: 5’-ATTTACATACAATRAAATCACCTC-3’;
ACE2_RT-PCR_Fw: 5’-CTCATGAGGAGGTTTTACTC-3’,
ACE2_RT-PCR_Rv: 5’-CATACAATGAAATCACCTCAAGAG-3’;
ACE2_Nested_Fw: 5’-AATGGGGTTTTGGCGCTCAG-3’,
ACE2_Nested-PCR_Rv: 5’-CTCAAGAGGAAATAAAATAG-3’;
ACE2_Nested_2^nd^_Fw: 5’-CTCAGGGAAAGATGTCAGGC-3’,
ACE2_Nested_2^nd^_Rv: 5’-TAGATTTCTCTAAAACGAAG-3’.

### CYTB sequence amplification from bat organs

Genomic DNA was extracted from bat liver tissue using the DNeasy Blood & Tissue Kit (QIAGEN, Cat# 69504), according to the manufacturer’s instructions. PCR primers^47^ were used to amplify the mitochondrial CYTB genes with KOD One PCR Master Mix - Blue- (TOYOBO, Cat# KMM-101). Amplicons were resolved by agarose gel electrophoresis using SeaKem GTG Agarose (Lonza, Cat# 50070). Target bands (*R. affinis* ACE2 DNA) were excised, and DNA was recovered and purified with a gel extraction kit (Qiagen, Cat# 28704) and tested with NanoDrop spectrophotometer (Thermo Fisher Scientific).

The primers used for this section are as follows:

CYTB_PCR_Fw: 5’- ACAGGCTCAAACAACCCAAC-3’,
CYTB_PCR_Rv: 5’- TGGCCTCCAATTCAGGTTAG-3’.

### DNA amplicons sequencing and analysis

Both ACE2 and CYTB amplicons from each sample were subjected to Sanger sequencing (Eurofins). Resulting reads were assembled with SnapGene v8.1.1 (SnapGene software, www.snapgene.com) and examined in AliView v1.28^48^. For ACE2 sequences, after alignment using MUSCLE v3.8.155158^49^ with default options we defined a set of unique ACE2 amino acid genotypes. To minimize false positives, two ACE2 samples were randomly selected from each putative genotype and subjected to TOPO cloning and transformation (TOPO Cloning Kit, Thermo Fisher, Cat# K4500-01SC; Champion DH5α chemically competent cells, SMOBIO, Cat# CC5204). For each sample, eight colonies were screened by colony PCR, purified, and the corresponding amplicons were Sanger sequenced (Eurofins). Consensus sequences were assembled and compared in SnapGene v8.1.1, and genotypes failing verification were discarded. This process yielded 15 high-confidence, reproducible ACE2 genotypes. In addition, comparison with all available *R. affinis* ACE2 genotype sequences downloaded from NCBI GenBank identified two genotypes not recovered in our sampling. These two genotypes were reconstructed by site-directed overlap extension PCR. In total, 17 ACE2 genotypes were retained for subsequent construction of ACE2 stable expressing cell lines (see “**Generation of HOS-TMPRSS2 cells stably expressing *R. affinis* ACE2 proteins**” section).

The primers used for this section for ACE2 sequencing are as follows:

ACE2_Seq_1_Fw: 5’-ATGTCAGGCTCTTCCTGGCT-3’,
ACE2_Seq_2_Fw: 5’-GAGGTCGGCAAGCAGCTGAG-3’,
ACE2_Seq_3_Fw: 5’-TTCAGGATCAAGATGTGCAC-3’,
ACE2_Seq_4_Fw: 5’-GACCATTTTTGAATTCCAGTTTC-3’,
ACE2_Seq_5_Fw: 5’-GAAGCCAAGAATCTCCTTCA-3’,
ACE2_Seq_6_Rv: 5’-CACTCTGCTGAAGAATCTGC-3’,
eACE2_Seq_7_Fw: 5’-GCCTATGAATGGAATGACAA-3’.

### Molecular phylogenetic analysis

To assess the phylogenetic relatedness between all sarbecovirus RBDs of interests, we manually selected 35 representative sarbecovirus RBDs displaying diverse ACE2 usage in the tree from our previous study^29^. We curated a panel of sarbecovirus spike glycoproteins that have been reported to utilize at least one ACE2 ortholog for cell entry in prior studies, with priority given to spikes originating from *R. affinis*. In addition, based on another report of isolation from *R. affinis*, we included the spike of virus Ra22DB163^38^. Finally, we included 36 sarbecoviruses spike in our study **(Table S3)**. The coding sequences corresponding to the RBDs of the 36 viruses were extracted and aligned with mafft v7.505^50^ using the --localpair option. A maximum likelihood (ML) tree was then reconstructed using IQ-TREE v2.3.6^51^ under the GTR+F+R10 substitution model with 10000 ultrafast bootstrap replicates^52^ **(Figure 2A)**.

For the *R. affinis* ACE2 trees presented in **Figures 1B and S2**, we extracted the ACE2 sequences for all *R. affinis* and constructed nucleotide sequence alignment using MUSCLE v3.8.155158^49^ under the default options. A single sequence of *Rhinolophus shameli* ACE2, the most closely related species with an available ACE2 sequence, was included in the alignment to be used as an outgroup. An ML tree was then reconstructed for this alignment using IQ-TREE web server (http://iqtree.cibiv.univie.ac.at/)^53^ under the best substitution model selected automatically by ModelFinder^54^ with 1000 ultrafast bootstrap replicates **(Figure 1B and S2A)**. The aligned ACE2 coding sequences were used as input for the GARD analysis (see “**Recombination analysis**” section). To account for recombination, the pre-aligned sequences were partitioned at the inferred breakpoint position into two segments: a left-hand (upstream) segment and a right-hand (downstream) segment. Phylogenetic trees were reconstructed independently for each segment, yielding a left-hand tree and a right-hand tree **(Figures S2B and S2C)**. All ACE2 phylogenies were rooted with the *R. shameli* ACE2 as the outgroup. Branch-specific nucleotide substitutions were inferred and annotated using TreeTime v0.11.3^55^.

### Recombination analysis

The aligned ACE2 sequences were used to perform recombination analysis with the Genetic Algorithm for Recombination Detection (GARD) method^33^, implemented in the DataMonkey online server (www.datamonkey.org/GARD)^56^, and run under the default options. GARD finished with a single breakpoint called at position 1685 of the ACE2 coding sequences, determined with a confidence value of 0.999.

### Selection analysis

Based on the recombination breakpoint identified, the ACE2 sequence alignment was split into two segments: the left-hand (upstream) and right-hand (downstream) sequences relative to the breakpoint. The split was made at the nearest codon site to the breakpoint. The dN/dS-based mixed effects model of evolution (MEME) method^34^ was used to test for site-specific selection separately for each of the two non-recombinant alignment partitions using the DataMonkey online server (www.datamonkey.org/MEME)^56^, under the default options. MEME found evidence of episodic positive/diversifying selection at 1 site with a p-value threshold of 0.05 within the left-hand partition and no evidence of selection within the right-hand ACE2 alignment partition.

### Plasmid construction

The plasmids expressing the human codon-optimized spike proteins of sarbecoviruses (summarized in **Table S3**) and human ACE2 protein were prepared in our previous study^29^ or synthesized by a gene synthesis service (Fasmac). Plasmids expressing the codon-optimized S proteins of Ra22DB163 and *R. affinis* ACE2 were generated by site-directed overlap extension PCR using the primers displayed below. The resulting PCR fragment was cloned into the KpnI/NotI site of backbone pCAGGS vector^57^ (for the S expression plasmids) or the BamHI/MluI site of pWPI-ACE2-zeo^45^ (for ACE2 expression plasmids) with 3×FLAG tag at the C terminus using In-Fusion HD Cloning Kit (Takara, Cat# Z9650N). Nucleotide sequences were determined by DNA sequencing services (Eurofins), and the sequence data were analyzed by SnapGene v8.1.1.

The primers used for this section are as follows:

pW_ACE2_Fw: 5’-CTAGCCTCGAGGTTTGGATCCGCCACCATGTCAGGCTCTTCC-3’,
pW_ACE2_Rv: 5’-AGTTTAAACACTAGTACGCGTCTACTTGTCATCGTCATCCTTGTAATCGATGTCATGATCTTTATAATCACCGTCATGGTCTTTGTAGTCAAACGAAGTCTGAAC-3’;
pC_Ra22DB163_Fw: 5’-CTATAGGGCGAATTGGGTACCATGATCTTTCTGATC-3’,
pC_Ra22DB163_Rv: 5’-AGCTCCACCGCGGTGGCGGCCGCTCAGGTGTAGTGCAG-3’.

### Generation of HOS-TMPRSS2 cells stably expressing *R. affinis* ACE2 proteins

HOS-TMPRSS2 cells stably expressing human ACE2 proteins were prepared as described in our previous study^29,41^. To prepare lentiviral vectors expressing *R. affinis* ACE2 in this study, LentiX-293T cells (500,000 cells) were co-transfected with 0.9 μg of psPAX2-IN/HiBiT, 0.2 μg of pCMV-VSV-G-RSV-Rev, and 0.9 μg of pWPI-ACE2-zeo using TransIT-293 (Takara, Cat# MIR2704) according to the manufacturer’s protocol. After 48 h of transfection, the supernatant including lentivector particles was harvested. HOS-TMPRSS2 cells (100,000 cells) were then transduced with the ACE2-expressing lentiviral vector. After 48 h post transduction, transduced cells were maintained for zeocin (50 μg/mL; Invivogen, Cat# ant-zn-1) and G418 (400 μg/mL; Nacarai Tesque, Cat# 09380-44) selections for 14 days.

### Western blotting

Samples were prepared for western blotting as previously described^41,58^. For the blot, 1 million of each HOS-ACE2/TMPRSS2 cells (see “**Generation of HOS-TMPRSS2 cells stably expressing *R. affinis* ACE2 proteins**” section) were harvested. The harvested cells were washed and lysed in RIPA buffer (50 mM Tris-HCl buffer [pH 7.6], 150 mM NaCl, 1% Nonidet P-40, 0.5% sodium deoxycholate, 0.1% SDS), protease inhibitor cocktail (Nacalai Tesque, Cat# 03969-21). The lysates were diluted with 2 × sample buffer [100 mM Tris-HCl (pH 6.8), 4% SDS, 12% β-mercaptoethanol, 20% glycerol, 0.05% bromophenol blue]. Both samples were boiled for 10 m. After cooling down, cell lysates were mixed with diluted sample buffer (ProteinSimple, Cat# 99351). Then, 5 × Fluorescent Master mix (ProteinSimple, Cat# PS-ST01EZ-8) was added at a ratio of 4:1. The Simple Western System was used for protein analysis. For protein detection, the following antibodies were used: horseradish peroxidase (HRP)-conjugated mouse anti-FLAG monoclonal antibody (clone M2, Sigma-Aldrich, Cat# A8592, 1:1,000), rabbit anti-β-actin monoclonal antibody (13E5, Cell Signaling, Cat# 4970, 1:1,000), and anti-rabbit secondary antibody (ProteinSimple, Cat# 042-206). Bands were visualized and analyzed using the capillary Western System Jess and Compass for Simple Western v6.1.0 (ProteinSimple).

### Pseudovirus infectivity assay

The pseudovirus assays presented in this study were performed as previously described^59–61^. Briefly, HIV-1-based, luciferase-expressing reporter viruses were pseudotyped with the spike proteins of sarbecoviruses and their derivatives. Lenti-293T cells (500,000 cells) were co-transfected with 0.8 μg psPAX2-IN/HiBiT^62^, 8 μg pWPI-Luc2^61^, and 0.4 μg plasmids expressing parental spike or its derivatives using TransIT-293 (Takara, Cat# MIR2704) according to the manufacturer’s protocol. Two days post-transfection, the culture supernatants were harvested, and the pseudoviruses were stored at −80°C until use. For pseudovirus infection, the amount of input virus was normalized to the HiBiT value measured by NanoGlo HiBiT lytic detection system (Promega, Cat# N3040) as previously described. In this system, HiBiT peptide is produced with HIV-1 integrase and forms NanoLuc luciferase with LgBiT, which is supplemented with substrates. In each pseudovirus particle, the detected HiBiT value is correlated with the amount of the pseudovirus capsid protein, HIV-1 p24 protein. Therefore, we calculated the amount of HIV-1 p24 capsid protein based on the HiBiT value measured, according to the previous paper. The amount of HIV-1 p24 antigen used in the assay was 4 ng (For all sarbecoviruses and related derivatives, except for Ra22DB163) or 24 ng (only for Ra22DB163). For target cells, the HOS-ACE2/TMPRSS2 cells (see ‘‘**Generation of HOS-TMPRSS2 cells stably expressing *R. affinis* ACE2 proteins**’’ section) were used. Two days post-infection, the infected cells were lysed with a Bright-Glo Luciferase Assay System (Promega, cat# E2620) and the luminescent signal was measured using a GloMax Explorer Multimode Microplate Reader (Promega).

### Association analysis between all 36 sarbecoviruses infection tropism and polymorphic sites in *R. affinis* ACE2

An ordinary least squares analysis (OLS) regression was performed to determine the potential association between amino acid residues at variable ACE2 sites. The OLS module of the statsmodels python package v0.14.2^63^ was used for performing the analysis. The regression was run separately for each virus with the log10-transformed infectivity values of each ACE2 genotype being the response variable and amino acid residues at all nine polymorphic sites (24, 34, 38, 49, 83, 185, 224, 300, 603) being the explanatory variables. To account for multiple testing, all resulting p-values were adjusted using a Bonferroni correction with alpha of 0.05. Data formatting, calculations, and numerical transformations were performed using the pandas python package v2.2.2^64^.

### Ra22QT77 mRNA vaccine production and mouse immunization

For mRNA vaccine preparation, the 5′ untranslated region, coding sequence and 3′ untranslated region were cloned sequentially downstream of a T7 promoter in an empty PSP73 plasmid. The plasmid was linearized by double digestion and used as the template for T7 RNA polymerase–driven in vitro transcription (Vazyme, Cat# DD4201) to produce mRNA encoding the Ra22QT77 S6P protein (stabilizing substitutions: F799P, A874P, A881P, A924P, K968P, V969P; amino-acid numbering corresponds to the Ra22QT77 spike). After transcription, reactions were treated with DNase I to remove template DNA and purified using VAHTS RNA Clean Beads (Vazyme, Cat# N412-02). A Cap-1 structure was added using Vaccinia Capping Enzyme (Vazyme, Cat# DD4109) followed by 2′-O-methylation with mRNA cap 2′-O-methyltransferase (Vazyme, Cat# DD4110), and the mRNA was cleaned again with magnetic beads. Poly(A) tails were added enzymatically with Escherichia coli Poly(A) Polymerase (Vazyme, Cat# N4111-02) and subjected to a final bead purification.

The mRNA was encapsulated in a functionalized lipid nanoparticle as described previously^65^. Briefly, ionizable lipid, DSPC, cholesterol and PEG2000-DMG were dissolved in ethanol at a molar ratio of 50:10:38.5:1.5. The mRNA was diluted in RNase-free 50 mM citrate buffer (pH 4.0) and mixed with the lipid solution at a lipid:mRNA weight ratio of 6:1. Aqueous and ethanol streams were combined using a microfluidic mixer at a 3:1 volume ratio. Formed LNPs were dialyzed overnight and stored at 2–8°C for up to one week to preserve component stability. Particle size, size distribution, encapsulation efficiency and mRNA concentration were measured; typical encapsulation efficiencies ranged from 90% to 99%.

Ten female BALB/c mice (6–8 weeks old) were used; no randomization or blinding was performed. Animals were maintained on a 12 h light/12 h dark cycle at 20–26 °C and 30–70% relative humidity. Each mouse received three intramuscular doses of Ra22QT77 LNP-mRNA (10µg per dose). The interval between dose 1 and dose 2 was 14 days, and the interval between dose 2 and dose 3 was 47 days. Plasma samples were collected 14 days after the second dose and 6 days after the third dose. On the day of dosing, LNP-mRNA vials were thawed on ice and diluted in sterile 1× PBS to the target concentration. Mice were injected intramuscularly into the quadriceps with 100 µL of vaccine using a 29-gauge insulin syringe; the needle was held in place for 3–5 s to minimize backflow.

### Neutralization assay

Pseudovirus neutralization assays against mouse vaccine sera were performed as previously described^60,66,67^. Briefly, the Ra22QT77 with other derivatives (pseudoviruses, counting ∼100,000 relative light units) were incubated with serially diluted (120-fold to 87,480-fold dilution at the final concentration) heat-inactivated sera at 37 °C for 1 hour. Pseudoviruses without sera were included as controls. Then, a 20 µl mixture of pseudovirus and serum/antibody was inoculated into HOS-ACE2/TMPRSS2 cells (V-Ra-10, 10,000 cells/100 µl) in a 96-well white plate. Two days post-infection, the infected cells were lysed with a Bright-Glo luciferase assay system (Promega, Cat# E2620) and the luminescent signal was measured using a GloMax explorer multimode microplate reader 3500 (Promega). The assay of each serum sample was performed in triplicate, and the 50% neutralization titre (NT50) or the 50% inhibitory concentration (IC50) was calculated using Prism 10 software v10.3.1 (GraphPad Software).

### Recombinant protein for structure analysis

The Ra22QT77 Ectodomain and *R. affinis* ACE2 sequences, each containing a C-terminal 6×His-tag, were independently subcloned into pHL-sec vectors (Addgene, Cat# 99845). Both proteins were co-expressed and secreted by HEK293 GnTI^ˉ^ cells. The cells were incubated at 37°C for 18 hours, and 10 mM sodium butyrate was added to induce expression. The cells were removed after 6 days at 30°C by centrifuging at 1,000 x g for 5 min, and the supernatant containing the secreted material was mixed with 20 mM Tris-HCl (pH 8.0) and 200 mM NaCl. The supernatant was incubation with Pierce High Capacity Ni-IMAC resin (Thermo Fisher Scientific, Cat# A50587) at 4 °C for 1.5 hours. The recovered resin was further washed with 15 column volumes of buffer containing 20 mM Tris-HCl (pH 8.0), and 500 mM NaCl, 20 mM imidazole. Proteins were then eluted with 20 mM Tris-HCl (pH 8.0), 150 mM NaCl, and 400 mM imidazole. The eluted fractions were concentrated and loaded onto a Superose 6 Increase 10/300 GL size exclusion column (Cytiva, Cat# 29091596) equilibrated with buffer containing 20 mM Tris-HCl (pH 8.0) and 150 mM NaCl.

### Cryo-EM sample preparation and data collection

The purified proteins were concentrated and added n-Octyl-β-D-glucoside (OG) (Nacarai Tesque, Cat# 25535-82) to a final concentration of 0.1%. The purified and concentrated complexes (3.4 mg/mL, 3 µL) were applied on the glow-discharged Quantifoil Au grids (R1.2/1.3, 300 mesh) (Quantifoil Micro Tools GmbH) and plunged into the liquid ethane using a Vitrobot Mark IV (FEI). Micrographs were collected using a 300 kV-Titan Krios G3i microscope (Thermo Fisher Scientific) equipped with a BioQuantum K3 imaging filter and a K3 direct electron detector (Gatan). In total, 7,309 movies were acquired using EPU software with a calibrated pixel size of 0.83 Å/pix and a defocus range of −0.8 to −1.6 μm.

### Cryo-EM image processing, model building, and model refinement

Data processing was performed using CryoSPARC v4.6.0^68^. Dose-fractionated movies were aligned using Patch Motion Correction and contrast transfer functions (CTF) were estimated using Patch CTF Estimation. Detailed data processing is shown in **Figure S4**. The density map was sharpened using Local Filtering. The quality of the density maps was sufficient to build the model manually using COOT for Windows version 0.9.8.93^69^. The model building was facilitated by ModelAngelo^70^. The model was then refined using phenix.real_space_refine version 1.21^71,72^.

### Predicted protein structure model

For **Figure 4C**, the RBD-only model of Ra22QT77 ins derivative was predicted using the AlphaFold3 online server^73^ under default options. The predicted RBD structure was aligned to Ra22QT77 cryo-EM RBD using matchmaker in Chimera X version 1.6.70^74^. The crystal co-structure of Ra22QT77 RBD and *R. affinis* ACE2 **(Figure 4D)** was used to set a consistent viewing orientation for **Figure 4C**. All protein structures were inspected and visualized using Chimera X version 1.6.70^74^.

### Mutation naming consistency in Ra22QT77

The naming of Ra22QT77, Ra22QT77 4sub, Ra22QT77 ins, and Ra22QT77 ins 4sub follows the conventions established in our previous study^29^. In this study, the four point mutants of Ra22QT77 are numbered consistently with the Ra22QT77 spike sequence and cryo-EM numbering. Our previous study^29^ used numbering corresponding to SARS-CoV-2 spike position numbering for the same four substitutions. For clarity we provide the corresponding substitutions here: SARS-CoV-2 (previous study) T346R, P486F, D487N, and Y496G; Ra22QT77 (this study) T328R, P468F, D469N, and Y478G respectively.

### Quantification and statistical analysis

In the pseudovirus assays, the infectivity was quantified using a GloMax Explorer Multimode Microplate Reader (Promega). All pseudovirus assays were performed in three biological replicates. Each 96-well plate contained an internal control group (HOS-TMPRSS2 cell line with human ACE2 receptor infected with SARS-CoV-2 spike). Raw signals were background-corrected (blank-subtracted) and normalized to the plate-specific means of the internal controls to yield relative infectivity. Three independent experiments were performed. Concordance across replicates was assessed by pairwise Pearson correlation on the normalized data; infectivity plots were generated from these normalized values accordingly. The infectivity value was shown as mean and SD in **Figures 3A** and **Data S1**. The NT50 of neutralization assay was shown as mean and SD and statistical significance was tested using a two-tailed Wilcoxon test in **Figure 3C**. The calculation of mean and SD and plot were finished using Prism 10 software v10.3.1 (GraphPad Software). All normalized infectivity data was used in **Figure 2A, 2B, 2D, 3A, 3B, and Data S1**.

## Supplementary information

**Figure S1.**
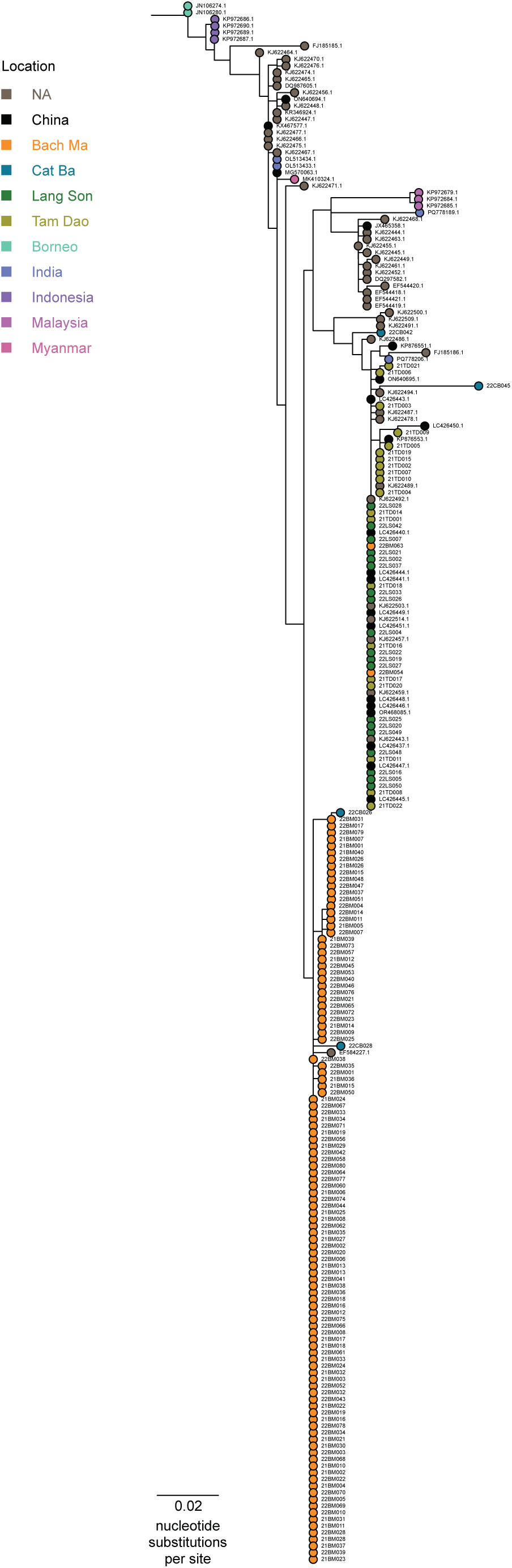
Maximum likelihood tree of *R. affinis* CYTB sequences. A total of 155 identified sequences and 79 sequences available in NCBI GenBank are included. TN+F+G4 is chosen as the best-fit substitution model. *R. shameli* ACE2 was used as the outgroup. Tips are coloured based on the locations where the sequences were sampled. The accession ID (from GenBank) and sample ID in this study are annotated at the tip label. Scale bars indicate nucleotide substitutions per site.

**Figure S2.**
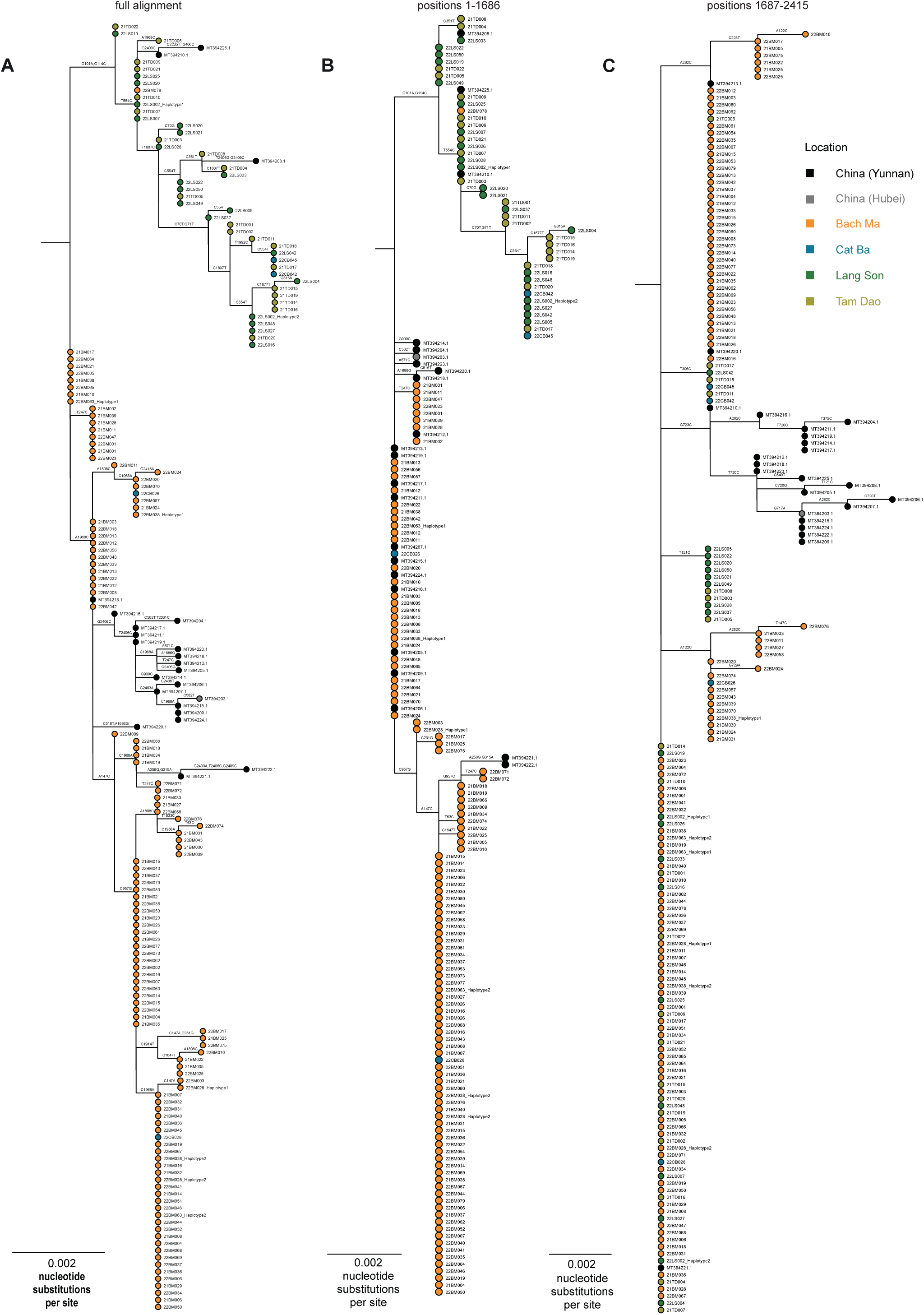
Maximum likelihood trees of the full-length and recombination breakpoint partitioned *R. affinis* ACE2 coding sequences. Phylogenetic trees were inferred from 182 sequences in total (159 study sequences plus 23 GenBank sequences). Trees are built from (A) the full alignment, (B) the left-hand fragment upstream of the inferred recombination breakpoint (positions 1-1686), and (C) the right-hand fragment downstream of the breakpoint (positions 1687-2415). All trees are rooted with *R. shameli*. Best-fit substitution models used were: (A) TIM3+F+I+G4, (B) K2P+R2, and (C) TPM3+R2. Tips are colored by sampling location, and tip labels include the GenBank accession (for references) and the sample ID from this study. Scale bars indicate nucleotide substitutions per site.

**Figure S3.**
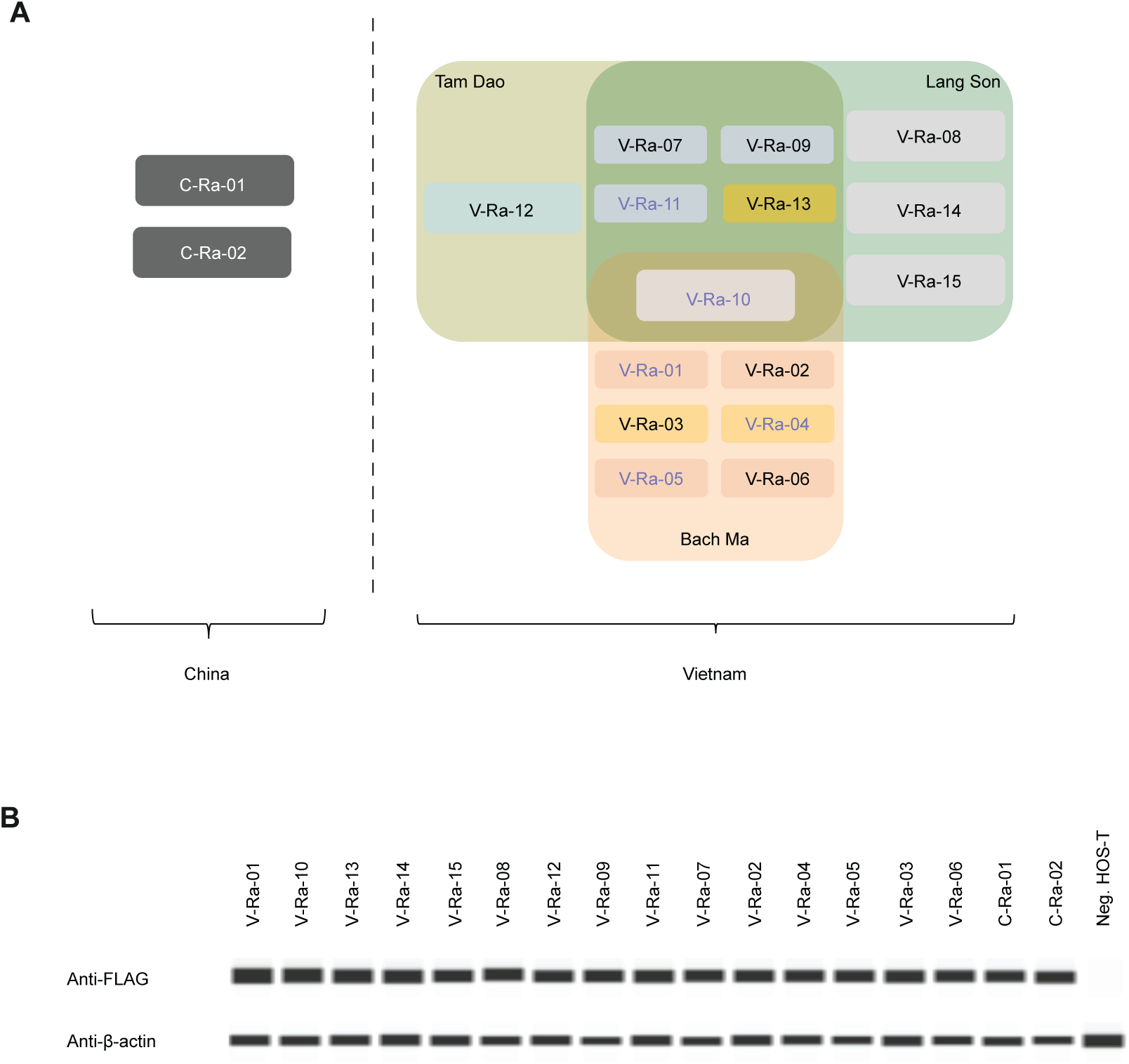
Geographical distribution of *Rhinolophus affinis* ACE2 genotypes and Western blotting test of HOS-A/T cell lines. (A) Venn diagram categorizing the 17 R. affinis ACE2 genotypes by their geographical locations. Three genotypes are highlighted in yellow to indicate their presence on Cat Ba Island, Vietnam, in addition to the annotated locations of Tam Dao, Lang Son, and Bach Ma. Five genotypes are marked with purple font to denote their previously reported presence in China, as described in earlier studies^24^. (B) ACE2 western blot for the 17 HOS-ACE2/TMPRSS2 cell lines used in this study. The original HOS-TMPRSS2 cell line is presented as a negative control. The FLAG protein is the target for testing with a FLAG gene specific antibody^29^. β-actin is blotted as the internal control for each cell line.

**Figure S4.**
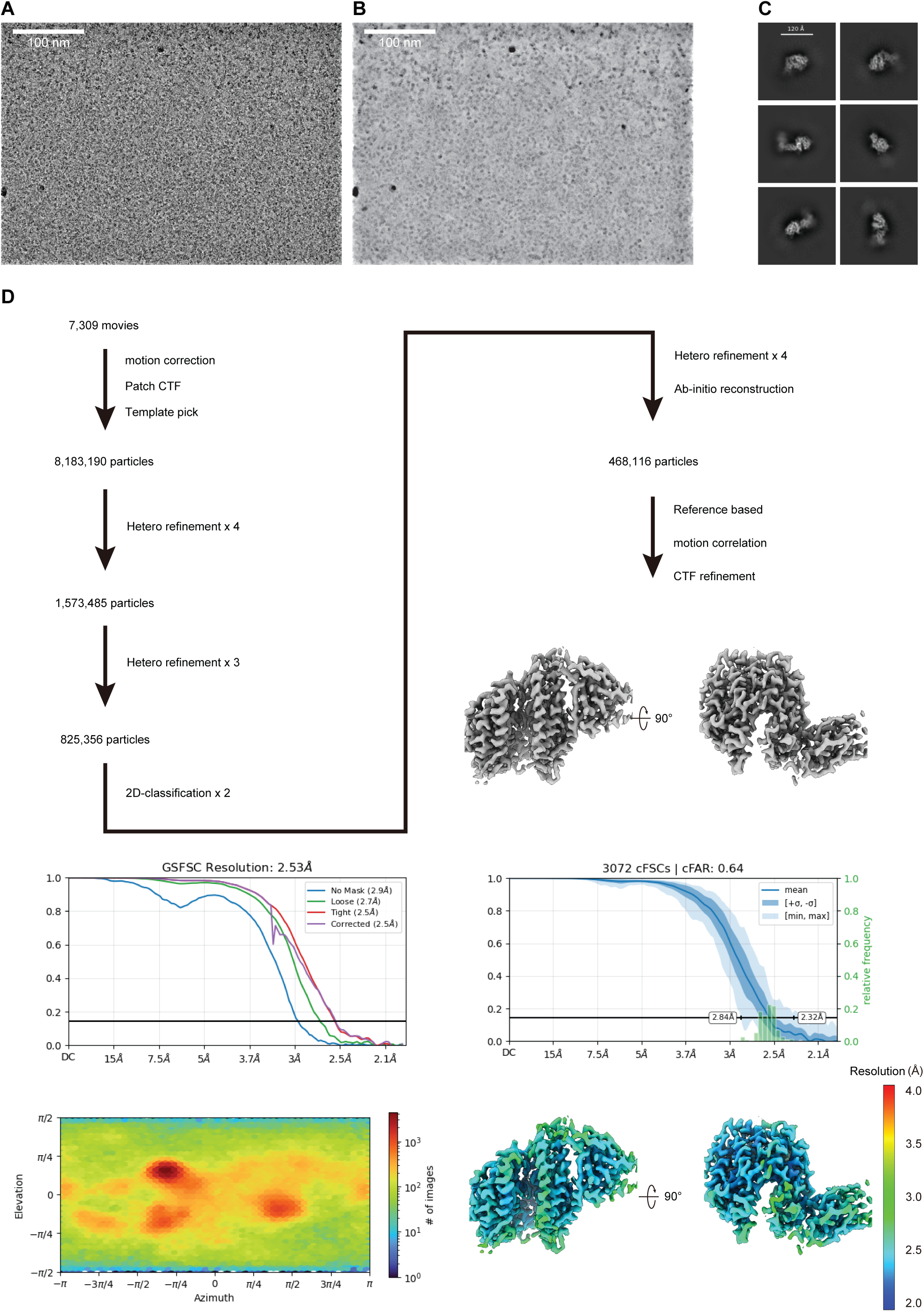
Single-particle cryo-electron microscopy analysis. (A)(B) A representative cryo-EM image (A), and a denoised image (B) of the Ra22QT77 RBD in complex with the *R. affinis* ACE2, recorded on a 300 kV Titan Krios with a K3 camera. (C) Representative 2D average classification classes. Scale bar, 90 Å. (D) Single-particle cryo-EM image processing workflow.

**Table S1. GPS coordinates and metadata for *R. affinis* sampling sites.**

Each row records location names, GPS coordinates of sampling site (latitude, longitude), sampling date, the number of samples captured, and the available Sanger sequences obtained. Date format: DD/MM/YYYY.

**Table S2. Metadata of *R. affinis* ACE2 sequences found and used in this study.**

Each row displays the accession number (NCBI GenBank) or sample ID (from our study), location where the sequence were derived from, genotype group name **(Figure 2A)**, lineage defined in our current study **(Figure 1B)**, and the name formerly used for publicly available ACE2 genotypes^24^.

**Table S3. Metadata of sarbecoviruses spike sequences used in this study.**

Each row shows accession number, virus name, host species, country where the virus was first found, the original source, and the sarbecovirus clade for each RBD as classified in a previous study^22^.

**Data S1. Detailed infectivity assay results.**

Infectivity plots of 17 *R. affinis* ACE2 genotypes infected with 36 pseudotyped sarbecovirus spikes. The infectivity data of each panel is normalized by an internal infection control (HOS-TMPRSS2 cell line with human ACE2 receptor infects with SARS-CoV-2 spike) and displayed as log value. Each bar plot included 12 replicates from 3 trials. The genotypes are ordered and categorized into 8 location-associated groups. TD, LS, CB, BM are abbreviations of Tam Dao, Lang Son, Cat Ba, and Bach Ma, respectively.

**Data S2. Cryo-EM data collection, refinement and validation statistics.**

Detailed information for the presented cryo-EM co-structure of the Ra22QT77 RBD bound to the *R. affinis* ACE2 (PDB: 9X23).

