## Supplementary figures and images for "Genetic diversity in horseshoe bat ACE2 and sarbecovirus spike proteins mutually shape one another"

### Supplementary Dataset 1

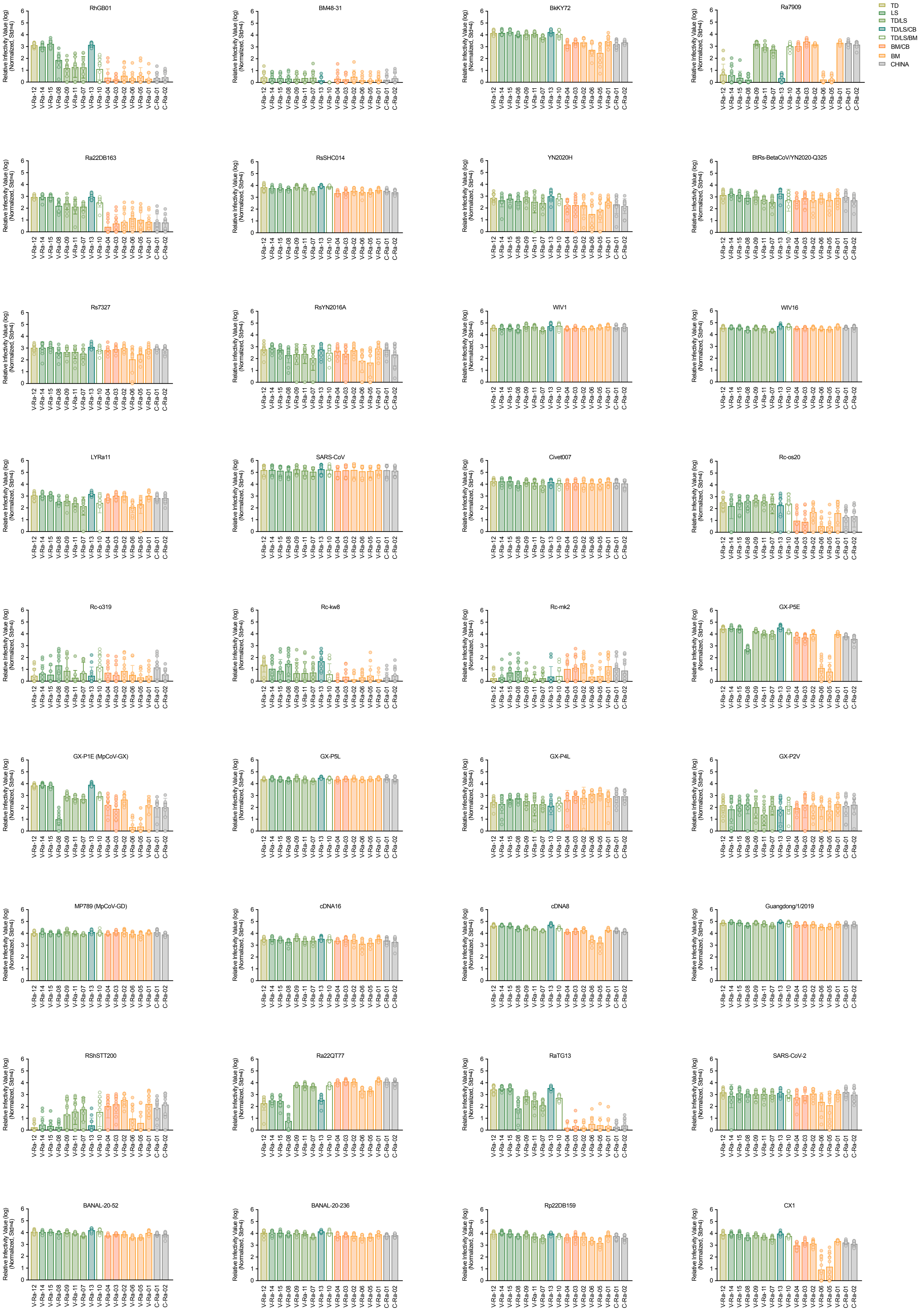

### Supplementary Figures

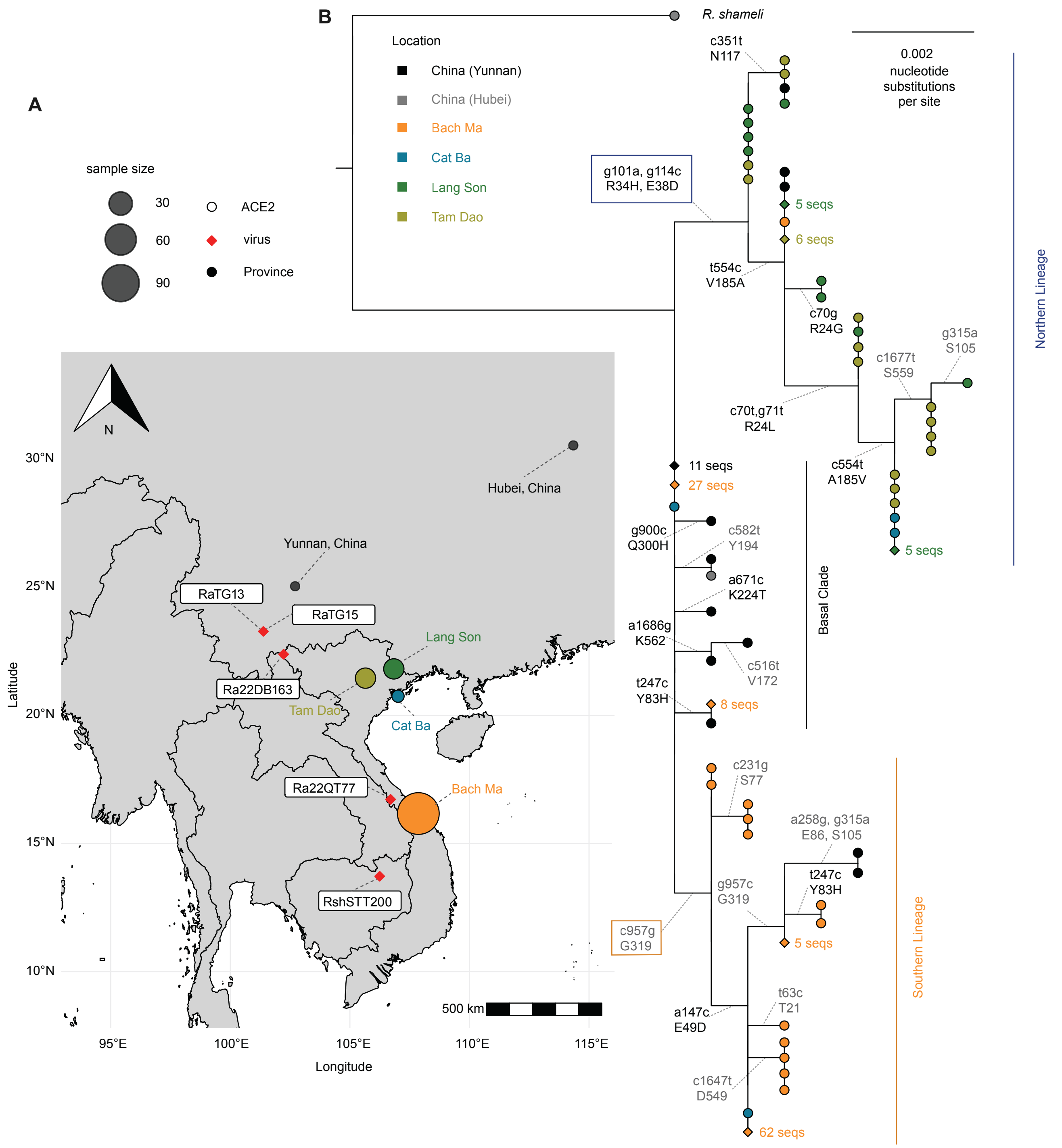

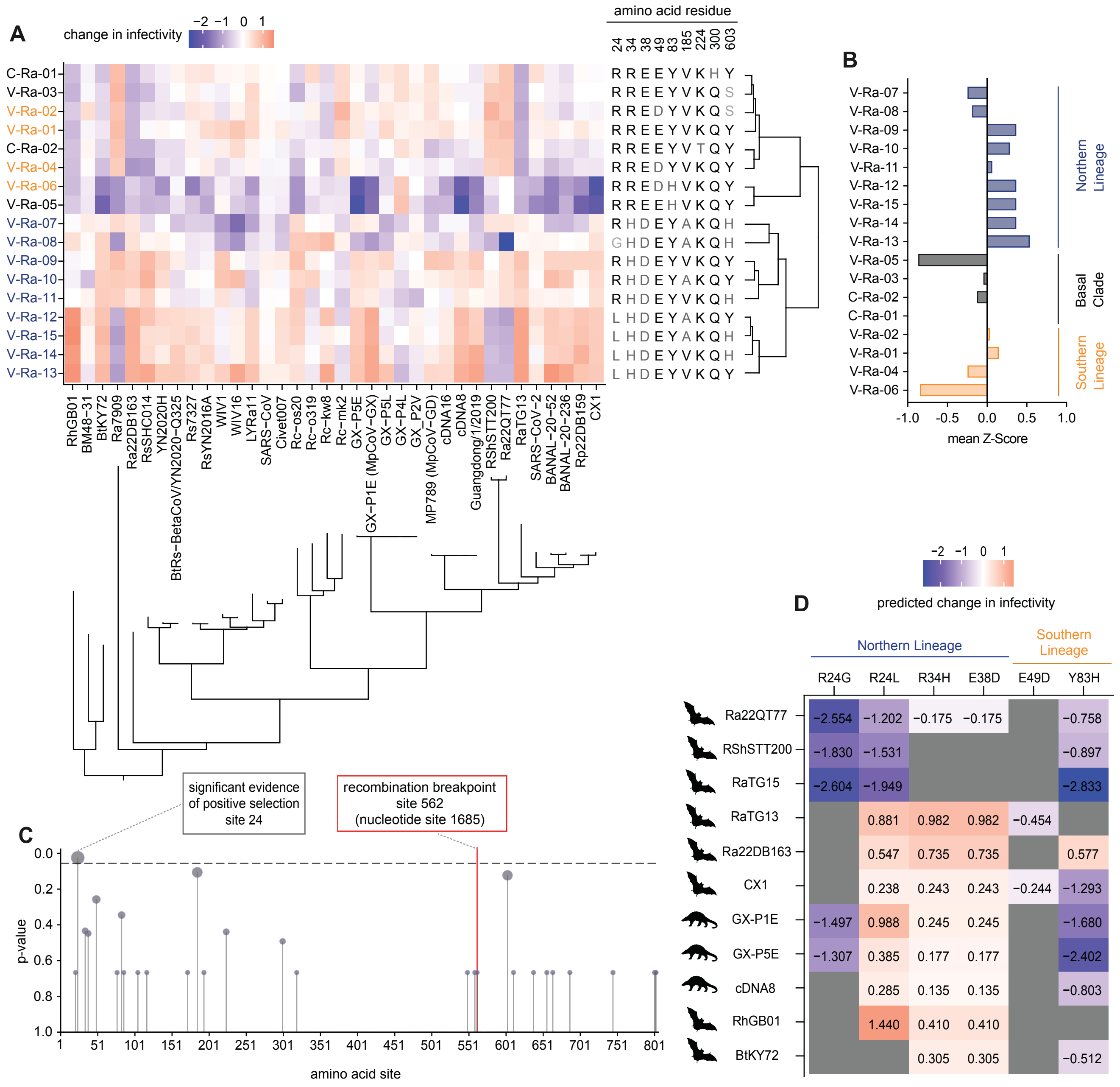

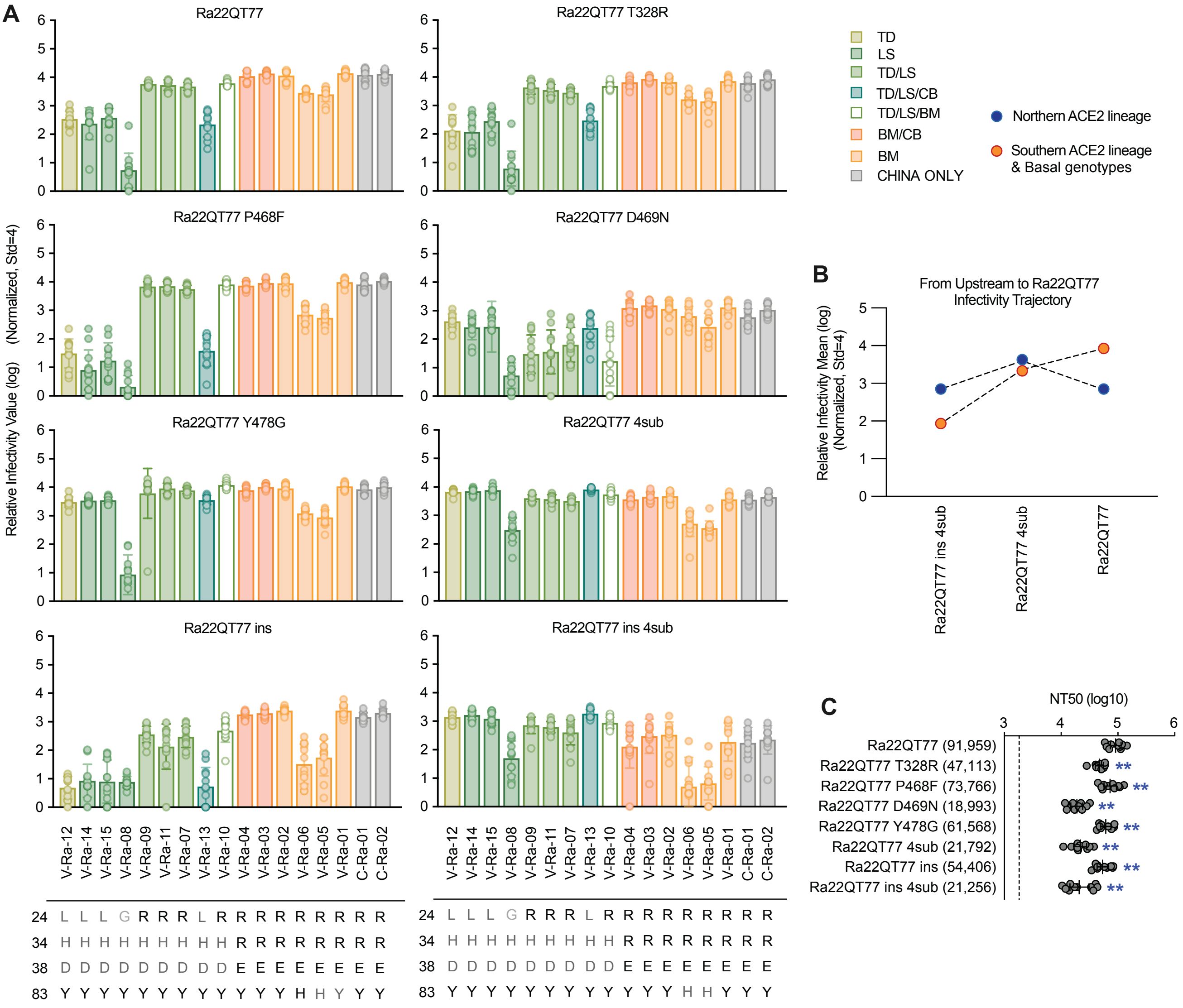

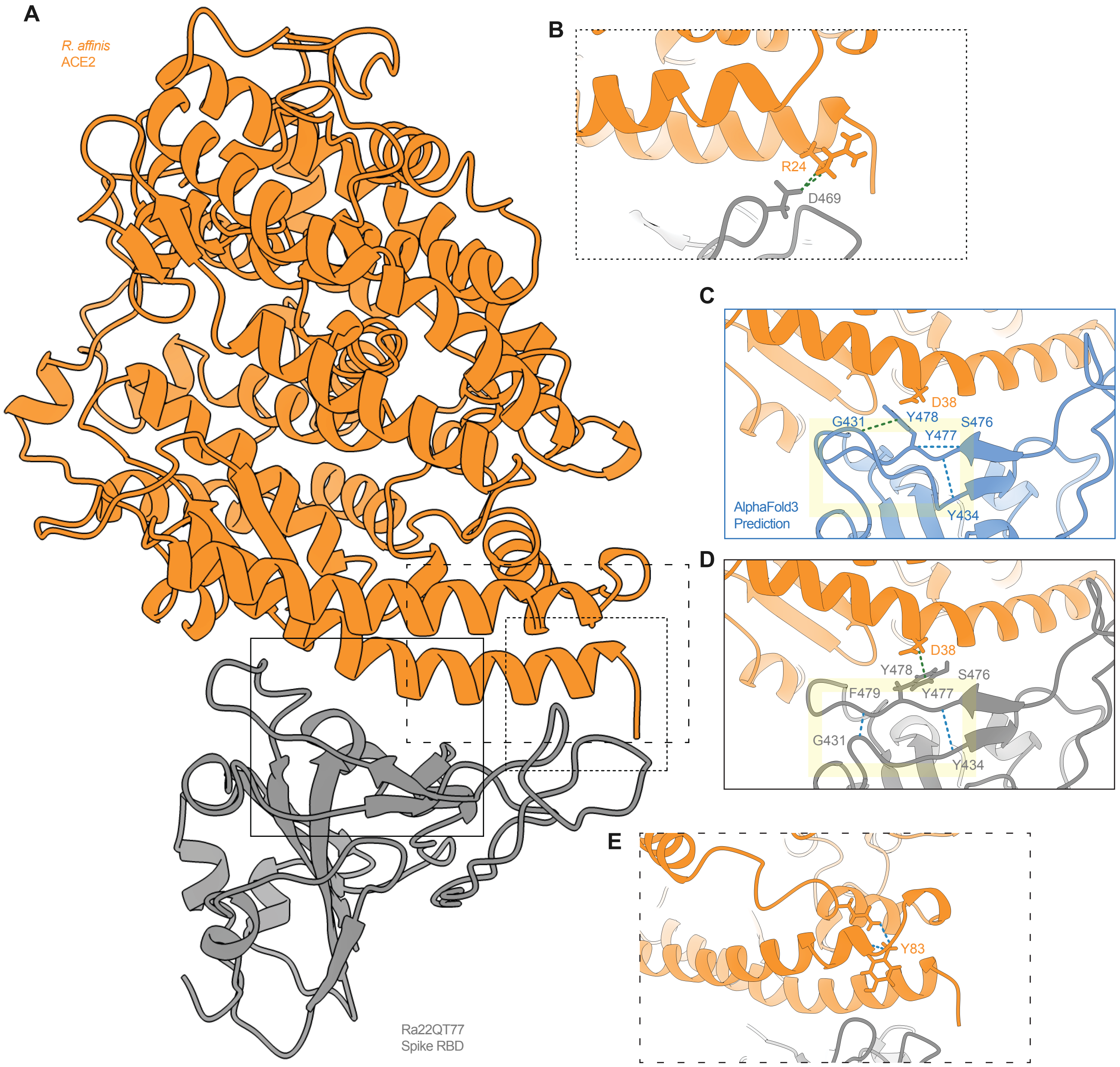

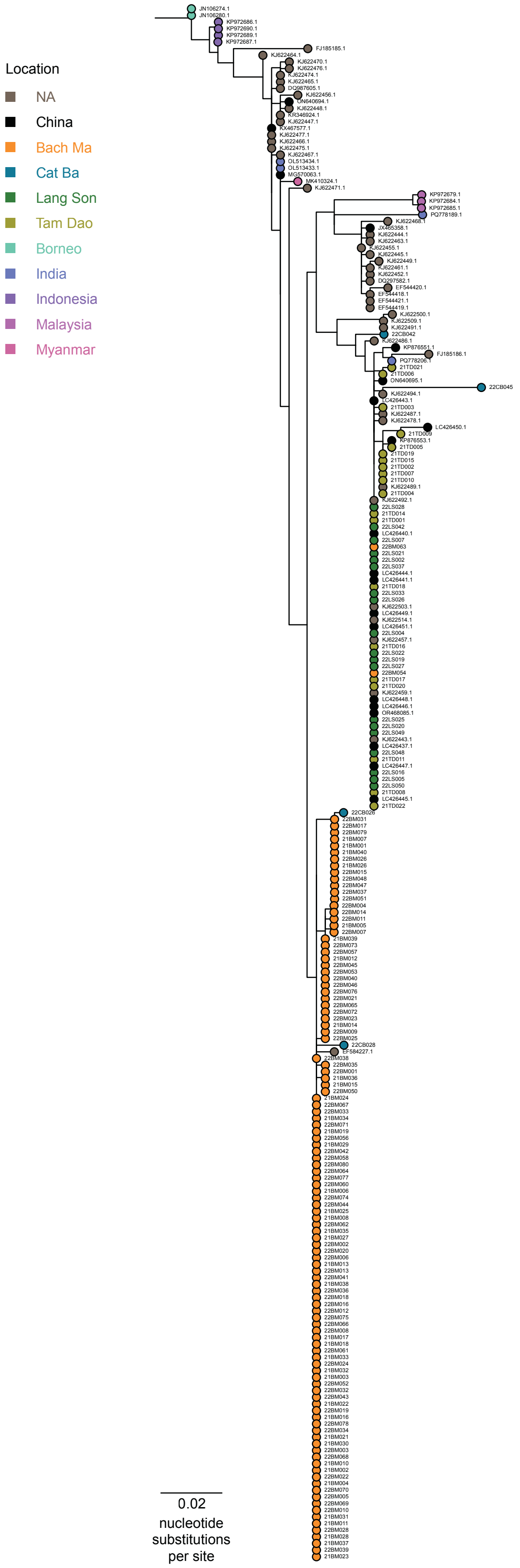

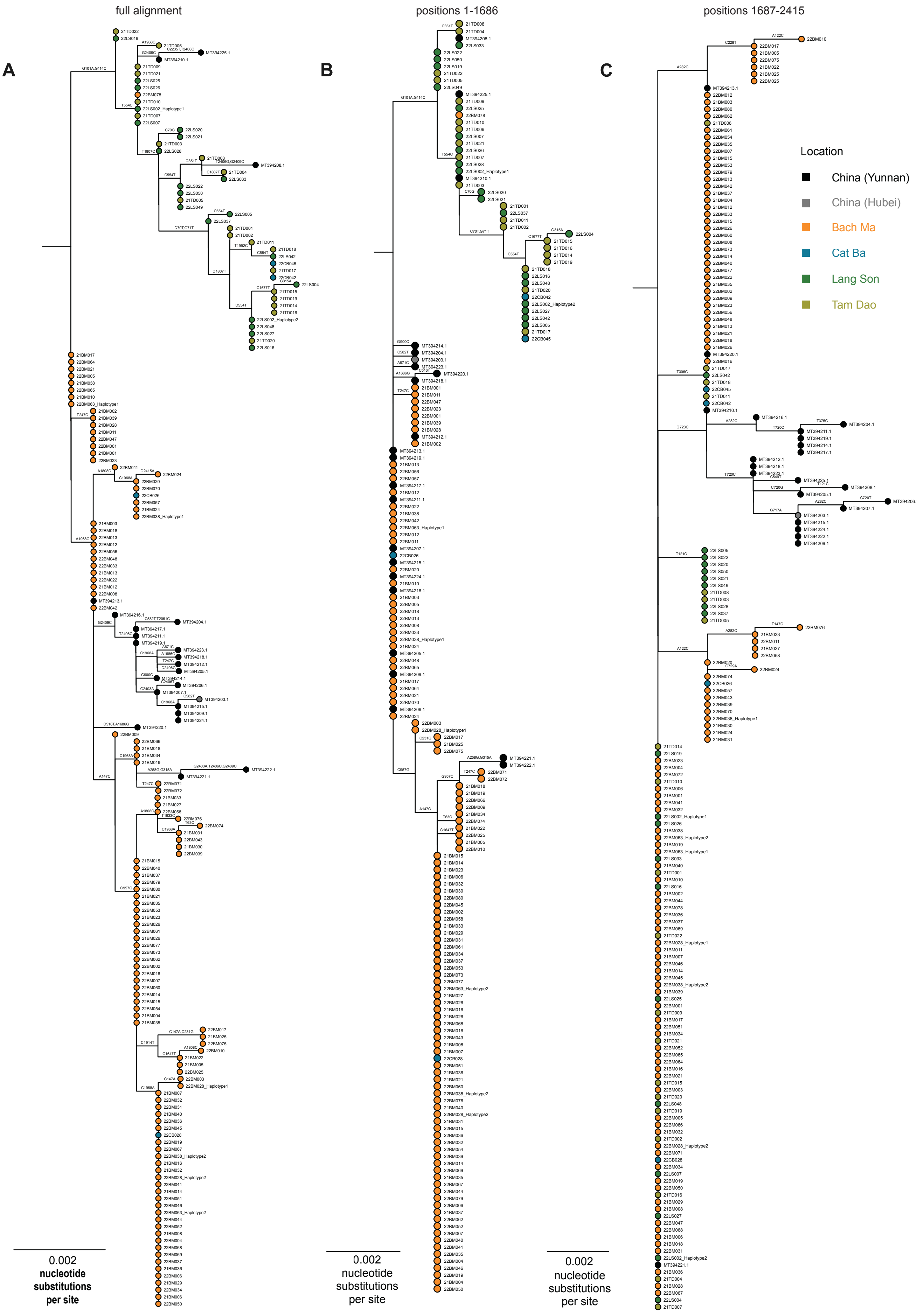

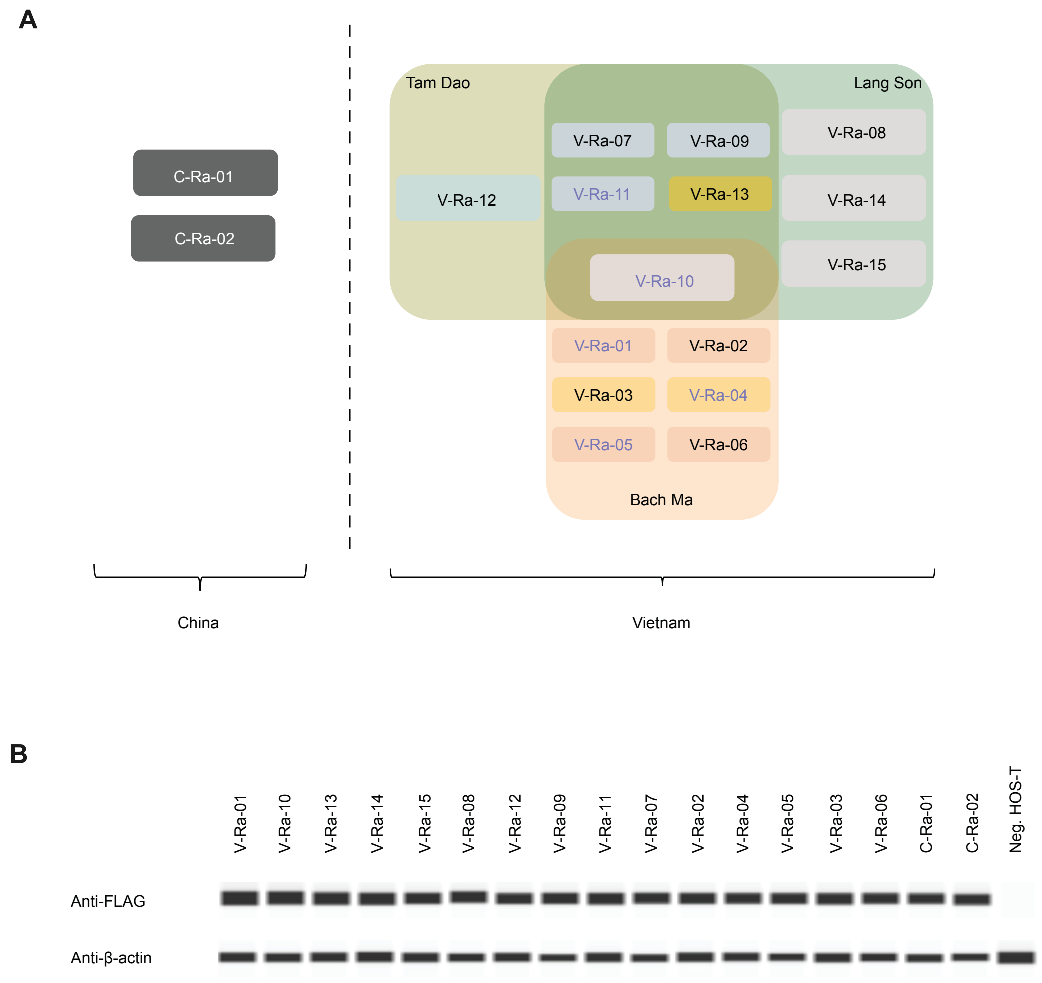

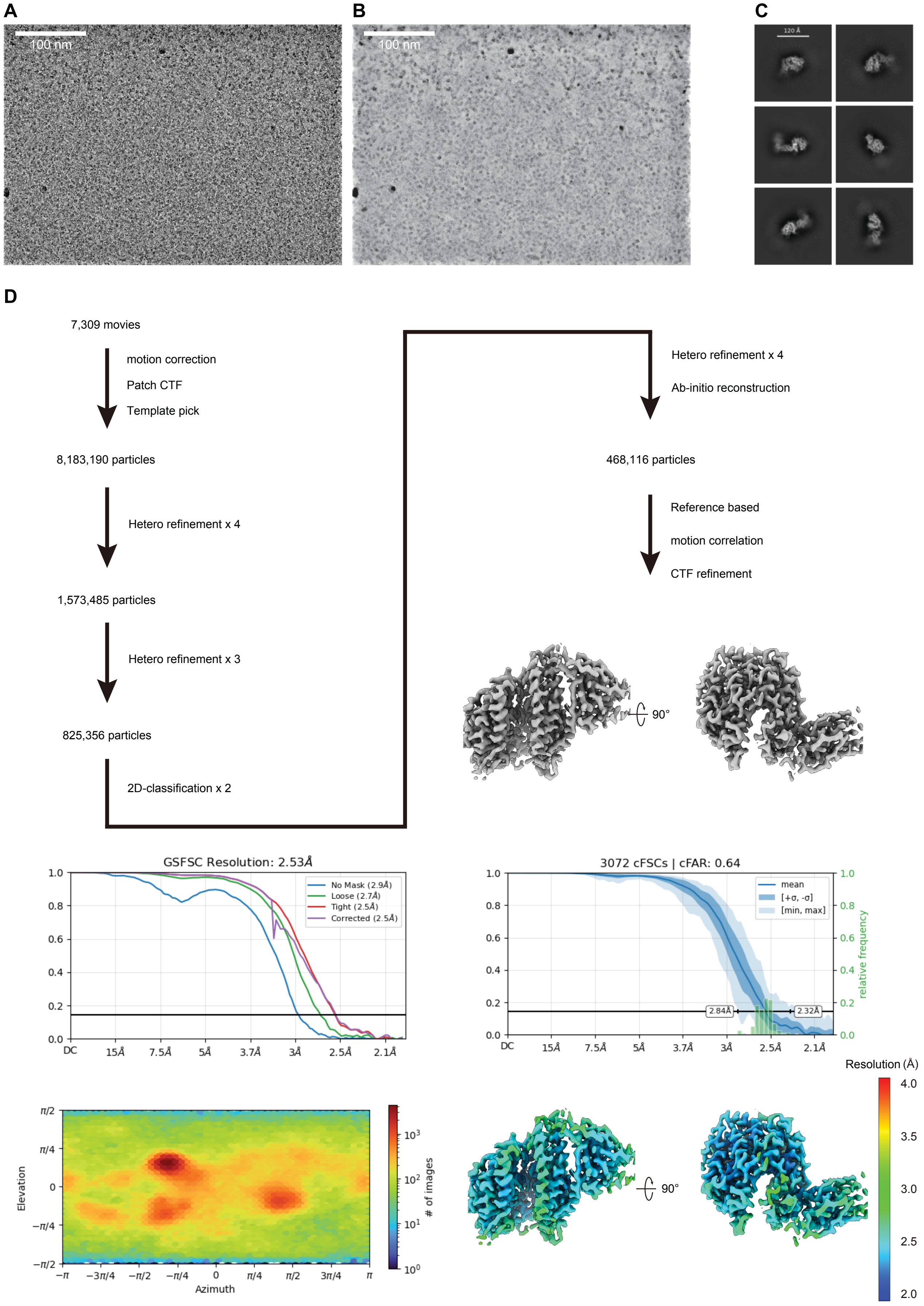
