## Supplementary Dataset 2 for "Genetic diversity in horseshoe bat ACE2 and sarbecovirus spike proteins mutually shape one another"

### Cryo-EM data collection, refinement and validation statistics

| | Ra22QT77 RBD in complex with Bat ACE2<br>(EMDB-66471)<br>(PDB 9X23) |
|---|---|
| **Data collection and processing** | |
| Magnification | x 105,000 |
| Voltage (kV) | 300 |
| Electron exposure (e–/Å$^2$) | 60 |
| Defocus range (μm) | -0.8－-1.6 |
| Pixel size (Å) | 0.83 |
| Symmetry imposed | C1 |
| Initial particle images (no.) | 8,183,190 |
| Final particle images (no.) | 468,116 |
| Map resolution (Å) | 2.53 |
| FSC threshold | 0.143 |
| Map resolution range (Å) | 2.198- 33.931 |
| **Refinement** | |
| Initial model used (PDB code) | N/A |
| Model resolution (Å) | 2.63 |
| FSC threshold | 0.5 |
| Model composition | |
| Non-hydrogen atoms | 6280 |
| Protein residues | 771 |
| Ligands | N/A |
| R.m.s. deviations | |
| Bond lengths (Å) | 0.004 (0) |
| Bond angles (°) | 0.615 (0) |
| Validation | |
| MolProbity score | 1.14 |
| Clashscore | 2.36 |
| Poor rotamers (%) | 0.29 |
| Ramachandran plot | |
| Favored (%) | 97.39 |
| Allowed (%) | 2.61 |
| Disallowed (%) | 0.00 |
